# Transposable elements drive intron gain in diverse eukaryotes

**DOI:** 10.1101/2022.06.06.494994

**Authors:** Landen Gozashti, Scott W. Roy, Bryan Thornlow, Alexander Kramer, Manuel Ares, Russell Corbett-Detig

## Abstract

There is massive variation in intron numbers across eukaryotic genomes, yet the major drivers of intron content during evolution remain elusive. Rapid intron loss and gain in some lineages contrasts with long term evolutionary stasis in others. Episodic intron gain could be explained by recently discovered specialized transposons called Introners, but so far introners are only known from a handful of species. Here, we performed a systematic search across 3,325 eukaryotic genomes and identified 27,563 Introner-derived introns in 175 genomes (5.2%). Species with introners span remarkable phylogenetic diversity, from animals to basal protists, representing lineages whose last common ancestor dates to over 1.7 billion years ago. Marine organisms were 6.5 times more likely to contain Introners than their terrestrial counterparts. Introners exhibit mechanistic diversity but most are consistent with DNA transposition, indicating that Introners have evolved convergently hundreds of times from autonomous transposable elements. Transposable elements and marine taxa are associated with high rates of horizontal gene transfer, suggesting that this combination of factors may explain the punctuated and biased diversity of species containing Introners. More generally our data suggest that Introners may explain the episodic nature of intron gain across the eukaryotic tree of life. These results illuminate the major source of ongoing intron creation in eukaryotic genomes.

## Introduction

The forces shaping intron-exon structures of eukaryotes remain among the longest-standing mysteries of molecular biology. Eukaryotic genomes contain from zero to hundreds of thousands of spliceosomal introns (*1*). Given the diverse roles of introns in gene expression and genome stability, from transcription enhancement to transcript surveillance to alternative splicing to R-loop avoidance (*2–5*), these differences may have important functional implications. Intron numbers per gene and per genome exhibit complex phylogenetic patterns, indicating massive recurrent changes in intron numbers through evolution, and comparative analyses attest to important roles for both intron deletion (loss) and creation (gain) (*1, 6, 7*). Despite decades of debate, no consensus has emerged as to either the proximal or ultimate explanations for these patterns.

Diverse molecular mechanisms of *de novo* intron creation are known but their relative contributions to genome evolution across the tree of life remain poorly understood. Proposed mechanisms of *de novo* intron creation include inexact double strand break repair (*8*), mitochondrial DNA insertion (*9*), internal gene duplication (*10*), and ‘intronization’ of exonic sequence (*11*). In addition to these *ad hoc* intron creation mechanisms, the intron-generating transposable elements (TEs) known as Introners represent a mechanism that could explain the high and episodic frequency and genome-wide scale of intron gains observed across eukaryotic lineages. These poorly understood TEs create introns *de novo* through insertion into exons (*12–17*). Introners have only been described in 5 eukaryotic lineages, and even among these cases, the precise molecular mechanisms remain obscure. Some Introner families show clear signatures of DNA TEs (*12, 15*) while others may be novel RNA-propagated elements (*14, 16*). More importantly, determining the extent to which introners are a primary source of ongoing intron gain is essential for interpreting the evolution of genome structure and function, and requires a broad survey that spans the eukaryotic tree of life.

By performing a systematic search and in-depth analysis of intron gain across all available eukaryotic genomes, we identified primary shared drivers of intron gain in diverse eukaryotic lineages. Our search identified 27,563 Introner-derived introns from 548 distinct families, with Introners found in 175/3325 (5.2%) of studied genomes. Introner-containing species span remarkable phylogenetic diversity, from copepods to poorly understood basal protists, representing lineages whose last common ancestor dates back to ~1.7 billion years ago (*18*). Unexpectedly, marine organisms were 6.5 times more likely to contain Introners, and 74% of Introner-containing marine genomes harbored multiple distinct Introner families. Overrepresentation in marine organisms could reflect higher rates of lateral gene transfer. While we find that Introners are efficiently spliced, preferential presence in lowly-expressed genes suggests that new insertions are costly. Most Introner families exhibit one or more signatures of DNA-based propagation. Our study indicates that susceptibility to acquire weakly deleterious Introners by lateral gene transfer might play the central role in a taxon’s tendency to gain introns.

## Results and discussion

### Introners are widespread across eukaryotes

Our survey across all available eukaryotic genomes indicates that introners are abundant in diverse lineages. To search for Introners, we developed a pipeline to systematically identify groups of introns with similar sequences, for which the region of sequence similarity extends to near the splice boundary at both ends. This approach allows flexibility in identifying introns created by TE insertions through potentially complex mechanisms while excluding most cases where inter-intron similarities reflect secondary insertion of TEs or evolution of microsatellites within pre-existing introns. We then applied this pipeline to 2,805 genomes representing 1,700 species with available genome annotations in Genbank (Supplementary Table 2). After extensive quality control (see Methods), our search revealed sets of Introners in 48 species grouping into 8 distinct taxonomic groups representing 6 major eukaryotic groups (Figure 1).

**Figure 1:**
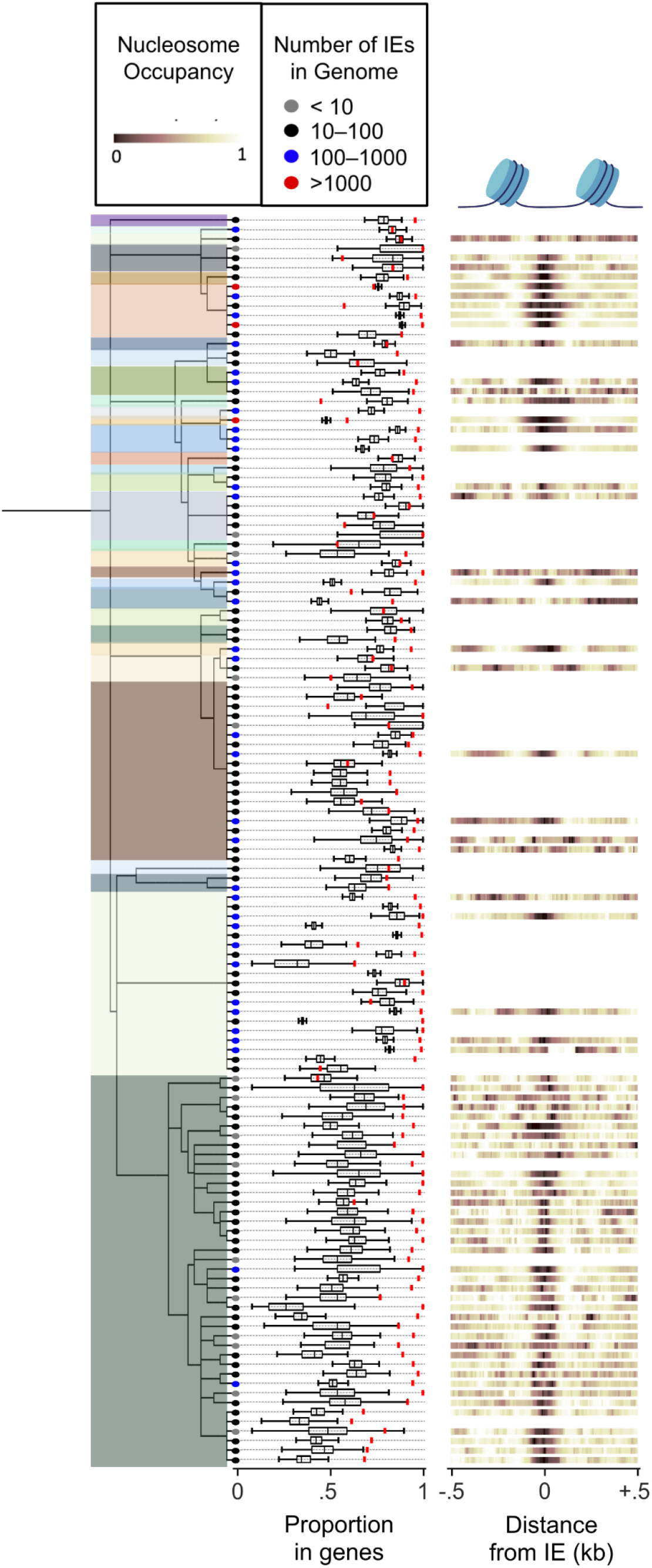
Diversity and characteristics of Introners across eukaryotes. Results are shown from 130 genomes representing 33 lineages with putatively independent acquisitions of Introners (different colors). Leaf tip colors indicate the total number of predicted Introners for each genome. Proportion in genes is shown by the red mark, which consistently exceeds the expected values as determined by randomization within each genome (black box plots; center line denotes median; box limits denote upper and lower quartiles; whiskers denote 1.5x interquartile range). Histone occupancy shows predicted histone occupancy of Introners, showing consistently reduced histone occupancy (dark) near Introner insertion sites. For genomes within the same genus, multiple genomes are shown only if the genomes have different complements of Introner families (see Methods).

### Introners are disproportionately common in marine lineages

Introners are disproportionately common in aquatic lineages, suggesting an important relationship between external environment and rates of introner evolution. To our surprise, 7/8 Introner-containing taxonomic groups (all except pezizomycotina fungi) inhabit aquatic environments. To further explore aquatic diversity, we analyzed 520 partial genomes representing 71 distinct genera (*19*) (Supplementary Table 3). This revealed 25 additional Introner-containing taxonomic groups, yielding a total of 33 separate taxonomic groups. Each presumably represents independent acquisition/evolution of Introners (Figure 1). Within this combined dataset, among 1,597 species for which aquatic/non-aquatic status was assignable, 17.0% of aquatic species (39/230) but only 2.6% of non-aquatic species (35/1367) exhibited at least one Introner family, confirming that marine habitat is significantly correlated with Introner presence (p < 10^-5^, two-sided Fisher’s exact test). Our results imply that aquatic environments are correlated with the evolution of introners and that aquatic environments may be an important driver of intron gain.

A test of environment association that accounts for phylogenetic relationship strengthens the conclusion that the genomes of organisms in aquatic environments are disproportionately likely to harbor introners. Because some introner-containing lineages are closely related, the apparent marginal correlation between aquatic environments and introner presence might be an idiosyncratic result of shared ancestry rather than an independently associated factor. We therefore retrieved a phylogeny of all eukaryotic species in our study and we found that a model where the rate of introner evolution depends on the environment is a significantly better fit than a model where the rate of introner gain is independent of the environment (p < 4.1×10^-4^ by a phylogenetic GLMM, Supplementary Table 4, See methods). This analysis therefore indicates that the strong correlation between environment and rates of introner gain is not strictly a product of shared ancestry, but rather may reflect an important determinant of rates of intron gain.

### Frequent convergent evolution of Introners from DNA transposons

Introner abundance varies substantially across introner-containing lineages and even between extremely closely related organisms. We identified 27,563 Introner-derived introns within 175 Introner-containing genomes, representing 548 putatively separate Introner families defined based on sequence similarity (Supplementary Table 6). Introner families exhibited substantial diversity in copy number per genome (5 to more than 2000), length (median length 30-654 nucleotides), GC content (20.9-83.3%) and percent of rare GC and GA 5’ splice sites (0-50.0% and 0-40.0%, respectively) (Supplementary Table 6). Genomes differed in the number of predicted Introner families, from one to 43 families (Figure 1, Supplementary Table 5). We also found large differences between closely-related organisms. For example, among three *Micromonas* species, an unknown species (isolate TARA_MED_95_MAG_00390) had no Introners, *M. commoda* had 44 total Introners, and *M. pusilla* had 3566 Introners with no sequence similarity to the *M. commoda* Introners. In *Florencialla,* some Introner families were found in all five isolates, and others in only a subset. We found similar patterns in the highest quality genomes, suggesting that data quality issues do not drive these results. Our findings suggest that introner presence and content is extremely evolutionarily labile, consistent with rapid changes in intron abundance observed across eukaryotic lineages.

Diverse molecular functions for intron gain suggest that many autonomous transposons convergently evolved into introners. Some previously characterized Introner families exhibit signatures of DNA cut-and-paste TEs while others lack such signatures (*12–14, 20*). Among Introner families for which TSD and/or TIR presence/absence could be ascertained (both characteristic of DNA TEs), 49/130 show evidence for TSDs and 72/177 show evidence for TIRs. Conversely, other families lack these features, including cases where Introners have 100% end-to-end sequence identity. For example, in *M. pusilla* both TSD/TIR families and non-TSD/non-TIR families are present (Supplementary Table 6). Introners also show remarkable diversity in mechanisms of splice-site recruitment. Among families with TSDs, we found a variety of orientations of splicing boundaries with respect to the TSDs. These included cases where the Introner carries the 3’ splice site and the 5’ splice site is recruited from the TSD (Figure 2B), cases in which the reverse is true (Figure 2C), or more complex cases in which either both splice sites are entirely or partially recruited from the TSD or from neighboring exonic sequence (Figure 2E). While most families followed previous reports in which insertion and splicing do not lead to a change in mRNA sequence length (*12, 13*), we also found cases where Introner insertion is associated with insertion or deletion of one or more neighboring codons (Figure 2F). This exceptional range of molecular mechanisms suggests that introners have independently evolved from diverse autonomous transposable elements and may explain recurrent bursts of intron gain across diverse eukaryotic lineages (*7, 21*).

**Figure 2:**
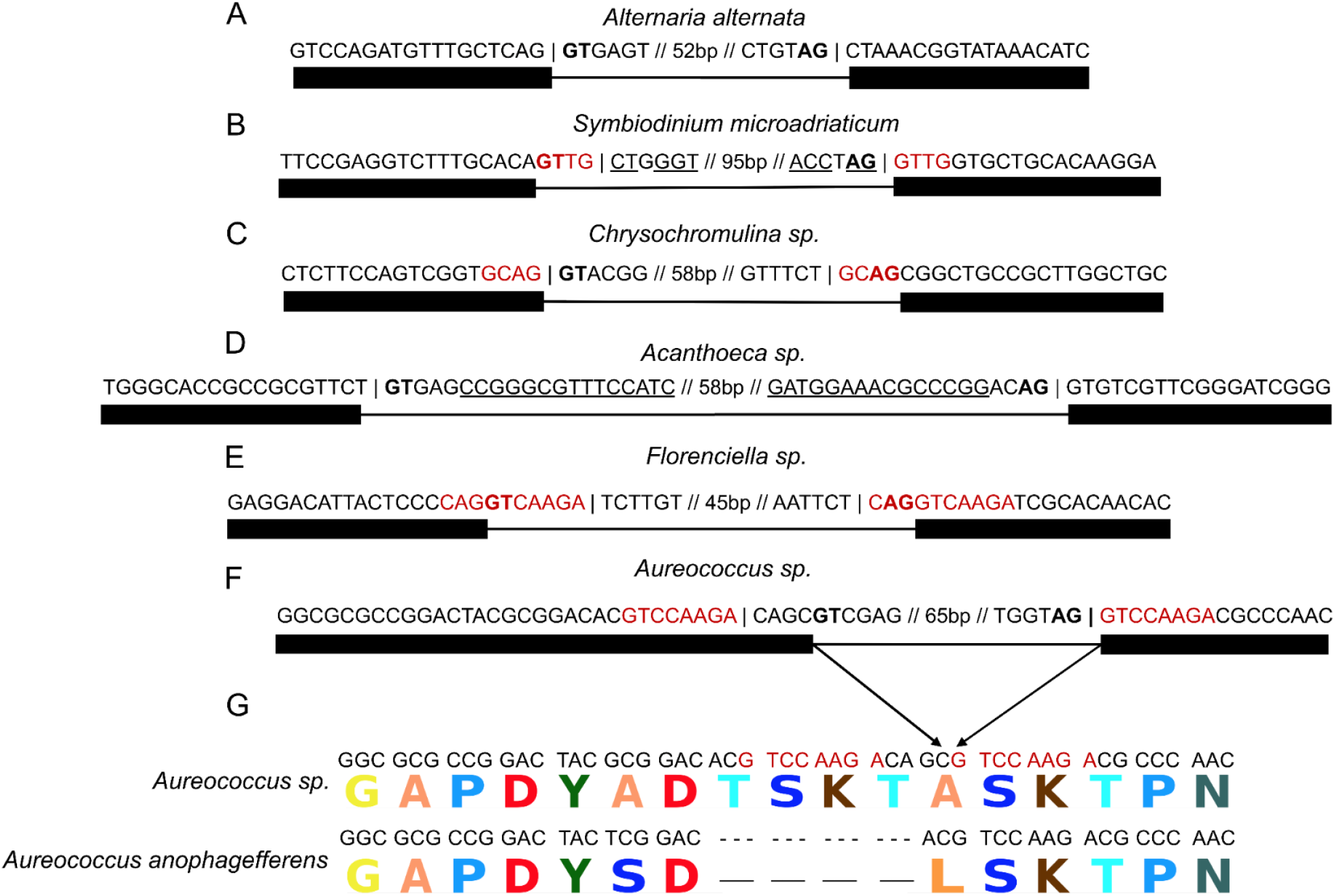
Examples of diverse Introner intron creation mechanisms. Splice sites are shown in bold, Introner boundaries are denoted by a vertical bar, and introns and exons are represented by lines and boxes respectively. (A) Introners in *Alternaria alternata* do not exhibit specific sequence features associated with known DNA transposition mechanisms and appear to replicate via direct insertion. (B) Introners in *Symbiodinium microadriaticum* show clear evidence of 4bp target site duplications (TSDs, shown in green) and terminal inverted repeats (TIRs, underlined), consistent with many known DNA transposons, carry their 3’ splice site, and co-opt their 5’ splice site from their TSDs upon insertion. (C) Introners in *Chrysochromulina sp.* show evidence of 4bp TSDs but no evidence of TIRs, carry their 5’ splice site, and co-opt their 3’ splice site. (D) Introners in *Acanthoeca sp.* show clear evidence of TIRs but no TSDs. (E) Introners in *Florenciella sp.* do not carry either splice site and instead co-opt both from their insertion site. (F) Introners in *Aureococcus sp.* carry both splice sites but add an extra 12bp into the transcript upon insertion (4bp from the Introner + 8bp from the TSD), (G) resulting in the addition of 4 amino acids to the respective protein when compared to an ortholog from a different isolate which lacks an Introner at that position.

### The majority of Introners may propagate via DNA-based mechanisms

Previous studies proposed different mechanisms for Introner mobilization. Some algal Introners appear to be MITE TEs based on observation of target site duplications (TSDs), terminal inverted repeats (TIRs), and biased insertion into nucleosome linker regions (*12*). In contrast, fungal Introners lack TSDs and TIRs and are highly biased towards gene regions (insertion into nucleosome linkers was not studied), and have been interpreted as novel RNA-based elements that propagate through reverse-splicing of RNA copies of spliced Introners (*14, 16*).

Although we observe exceptional molecular diversity (above), most Introner families exhibit at least one signature consistent with DNA transposition and ascomycetes are an outlier. Among 130 families for which both TSD and TIR presence/absence could confidently be determined, 59.2% have either TSDs, TIRs or both (78.5% when we exclude ascomycetes fungi, see below). Presence of separate DNA- and RNA-based families predicts that putative DNA-based signatures are positively associated with each other across families, and are negatively associated with the putative RNA-based signature of bias towards genes. However, TSDs are present in equal fractions of TIR-containing and non-TIR-containing non-ascomycetes families (51.7% (30/58) and 48.7% (19/40), respectively; p = 0.99 two-sided Fisher’s exact test), and we did not find an association between TSD or TIR presence and nucleosome linker bias (Supplementary Table 7). Furthermore, there is little association between TSD or TIR presence and bias towards insertions in genes (Supplementary Table 7, see Methods). We also found no difference when comparing families that differed in TSD/TIR presence when accounting for species, suggesting that correlations were not obscured by unaccounted for interspecific differences (Supplementary Table 7). Ascomycete introner families are an outlier, with no TIRs or TSDs. Together, our results suggest that most Introners propagate via DNA transposition.

### Introners show insertional preferences at various genomic scales

We next investigated the signatures of Introner insertion by studying Introner insertion positions at the level of genome region, nucleotide content, and chromatin structure. Introners are enriched in genes in 161/175 Introner-containing genomes (Figure 1, Supplementary Table 5). This pattern echoes some DNA elements, for example *piggyBac* elements in *Drosophila,* which also preferentially insert into coding regions (*22*). Overrepresentation of Introners within genes could also reflect an insertional bias towards GC-rich regions, since genes are typically more GC-rich than intergenic regions. GC-bias has also been previously reported for many DNA TEs (*23*). We find that Introner families enriched in genes also tend to show biased insertion into GC-rich motifs (P < 0.0001 binomial test, Supplementary Fig 1,5). One possibility is that TEs that exhibited a preexisting bias for insertion in GC-rich genic regions experience greater selection for efficient splicing to reduce gene disruption thereby creating new introners. Finally, some putatively DNA-based Introners in *M pusilla* and *Aureococcus anophagefferens* preferentially insert in nucleosomes linker regions (i.e., between nucleosomes (12)), similarly to many DNA transposons (*24*). Computationally predicted nucleosome occupancy profiles showed a bias towards linker region insertion for 78.8% (104/132) of Introner-containing genomes for which prediction was possible (Figure 1, Supplementary Fig 2-3, Supplementary Table 5-6).

### Negative selection shapes the distribution of Introner insertions

Introners exhibit a range of molecular phenotypes indicative of negative selection on the majority of new insertions. We find that observed Introners are generally more efficiently spliced than are other introns (p = 5.9×10^-3^, binomial test, Figure 3A-B). Because observed Introners may not reflect splicing of all new insertions it is likely that more deleterious insertions could be removed by selection and thus absent from sequenced genomes. This pattern suggests that high frequency Introners may have limited mis-splicing-related fitness costs due to negative selection purging Introners that are frequently misspliced. Similarly, we find that Introner insertions are biased towards lowly expressed genes relative to other introns (p = 2.4×10^-2^ sign test, Figure 3C), as could be expected if some Introner insertions impose transcription- or splicing-associated costs. This bias is unlikely to result from an insertion site preference given preferential Introner insertion into GC rich genes, which are typically more highly expressed (*25*). These data therefore suggest that negative selection impacts the range of introner insertions we observe.

**Figure 3:**
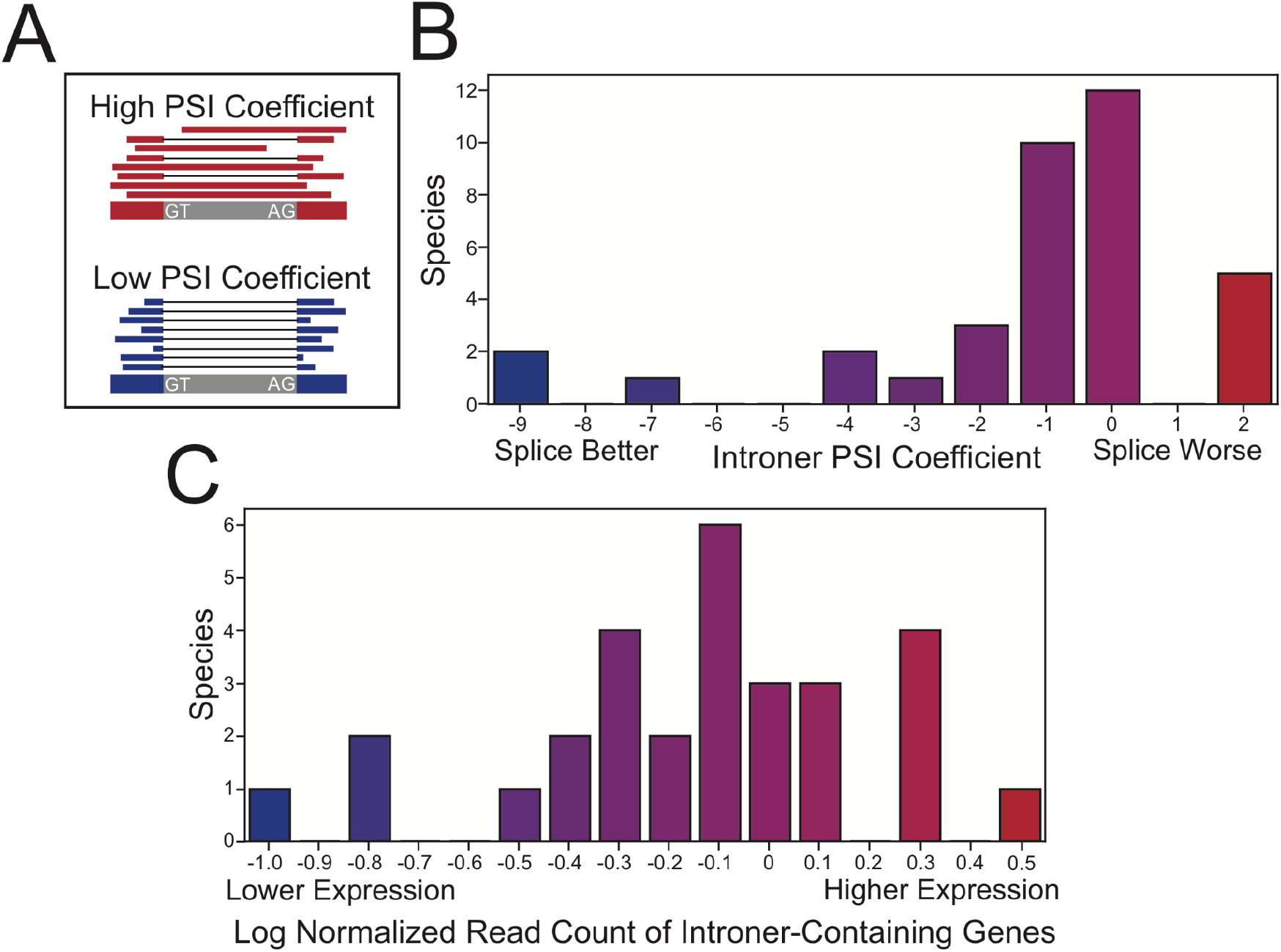
Relative to other introns, Introners are more efficiently spliced but less frequently found in highly expressed genes. (A) Explanation of PSI. Species with greater PSI coefficients (above 0, red) have Introners spliced in more frequently than other introns in the same genome. (B) Introners are more efficiently spliced than other introns in most species, as indicated by PSI coefficients less or equal to 0 for 31/36 species. (C) Introners are overrepresented in lowly expressed genes for most species. Relative log-normalized gene expression values for Introner-containing genes relative to other Introner-containing genes are shown.

### A new model for the evolutionary forces governing intron gain

Our results suggest that a range of previously proposed models for the major forces governing intron gain require reconsideration. We find no clear support for previous influential proposals that organismal complexity or small population size promotes intron gain (*26*). Most strikingly, among animals and land plants, two organismally complex groups with typically small effective population sizes, we find Introners in only two taxonomic groups (both animals), despite accounting for one-quarter (835/3325) of studied genomes (*p*<0.00001, two-sided Fisher’s exact test). This dearth of Introners is all the more striking given that these lineages are intron-rich, are generally more slowly-evolving at the sequence level which facilitates Introner discovery, and widely use introns in gene regulation (*1*). Nor do we see evidence for a pattern of Introner gain in species whose biology predicts small population size, such as parasites.

We propose that the distribution of introners reflects the propensity of lineages to acquire new genetic elements via horizontal gene transfer (HGT). All Introner-containing lineages except Ascomycetes and one species of blastocyst are marine organisms with marine organisms mostly represented by a remarkably diverse array of unicellular organisms. We propose that this pattern reflects greater rates of HGT for these species. Indeed, marine environments generally favor HGT (*27–30*), and marine unicellulars in particular have been shown to exhibit high rates of HGT possibly because they often live in dense microbial communities and require interactions with other species for their ecology (*31*). A variety of studies have detailed large-scale lateral gene transfer in marine protists, including several that have Introners (*31–34*). The only two non-marine Introner-containing lineages, ascomycetes and blastocysts have also been reported to have large amounts of HGT (*35, 36*). Furthermore, DNA transposons are apparently well adapted for HGT and have frequently made jumps between highly diverse lineages (*37*). The introduction of a new DNA transposon unfamiliar to a host might also intensify selection for the independent evolution of Introners. Newly acquired TEs often evade host-mediated TE silencing mechanisms and thereby have the freedom to mobilize at high frequencies, representing a major cost to host fitness (*37*). The ability of a TE to be spliced could alleviate some of these costs. Thus, HGT could not only explain the highly punctated pattern of Introner presence across distantly related taxa, but also favors the independent evolution of Introners in new lineages.

## Conclusion

The proximate and ultimate origins of introns remains a fundamental question in biology. Here, we demonstrate that Introners generate new introns on genomic scales in a remarkable diversity of eukaryotic lineages. Despite many similarities, the extensive molecular diversity that underlies Introner transposition reveals that a vast range of transposon families have independently evolved into Introners. In light of these findings, frequent horizontal transfer of TEs and the extreme marine biased distribution of species harboring introners, we propose that a crucial factor governing lineages’ tendency to gain introns over time is exposure to transfer of TEs from diverse unrelated eukaryotic organisms.

## Acknowledgements

The authors thank J. Collemare, A. van der Burght, and J. Logsdon for helpful discussions, as well as several community members who participated in a conversation about the nomenclature for Introners. We also thank Cedric Feschotte for helpful feedback on this manuscript.

## Author contributions

LG, RCD and SWR conceived and designed the study. LG, RCD and SWR developed the pipeline for Introner detection. LG and SWR wrote code for Introner detection, validated Introner candidates, and performed all analyses except the RNA analysis and prediction of candidate transposes which were performed by BT and AK respectively. LG and BT produced figures and tables and LG, RCD, and SWR wrote the manuscript. All authors analyzed and interpreted results. LG, RCD, SWR, and MA edited the manuscript.

## Methods

### Accessing genomic data

We performed our systematic search for Introners on all annotated genomes in Genbank (last accessed 9:24 AM 04/10/2020). We used ftp links available through a csv file downloadable from NCBI (Supplementary Table 1) to systematically access and download genomic data for our analyses. We filtered out genomes that lacked annotation files or for which annotations were dubious based on low gene number (Supplementary Table 2). Fasta files and genome annotations were downloaded for the Tara Oceans Eukaryotic Genomes project, from https://www.genoscope.cns.fr/tara/. Genomes without gff annotation files were filtered out, yielding a total of 520 genome assemblies.

### Identifying highly similar introns

To find candidate Introners, for each genome, we first extracted all annotated intron-exon structures, that is, genomic sequences corresponding to annotated protein-coding sequences spanning from translation start to stop codons, and with exon and intron sequences indicated by upper and lower case. We then extracted each intron along with up to 20 nucleotides of flanking sequence (requiring a minimum of 10 flanking exonic nucleotides). We then searched for introns with sequence similarity to each other. Candidate similar pairs were identified by an all-against-all blast (*38*) of introns-plus-flanking sequences within each species, with a minimum e-value of 10^-5^. Previous results and our preliminary findings reveal that there exist several reasons that two introns can have extensive sequence similarity other than creation by the same Introner family, therefore we performed several filtering steps to eliminate false positives.

We filtered for two alternative reasons for intron-intron sequence similarity. First, many introns with extensive sequence similarity to each other owe this similarity to secondary insertion of transposable elements or to microsatellites within the intron interior. Conversely, introns from paralogous genes or gene regions can be similar due to duplication of a longer region. These two cases share a frequent signature, namely that the region of intron-intron sequence does not correspond to (roughly) the whole of both introns, but instead is either a subportion of the intron (in the first case) or extends beyond the intron (in the second case). Therefore we required that the region sequence similarity between the two introns extend to near the exon-intron boundary. After iterative manual scrutiny, we chose to require that sequence similarity begin within a 15 base region spanning 5 exonic bases or 10 intronic bases, and that this be the case for both introns for both 5’ and 3’ intronic boundaries.

After initially requiring end-to-end blast hits, we learned that secondary indels including secondary TE insertion led to many false negatives. Consequently, we applied a different strategy where we judged similarity between each pair of introns based on pairwise similarity of the two ends, either similarity between corresponding ends (5’ with 5’, 3’ with 3’, as expected from same-orientation insertion) or opposing ends (5’ with 3’, 3’ with 5’, as in opposite-orientation insertion). Up to 100 intronic nucleotides were assessed for each end. We required that the similarities were in the expected orientation (i.e., the interiors of the two introns lining up together).

Recent paralogous gene duplications can result in sequence similarity between intron sequences. While the requirement that nucleotide-level sequence similarity begin an end near the intron-exon boundary removes most such cases, we also observed cases where rapid exonic evolution (or simple chance substitution) led to paralogous sequences being retained (the clearest case involved the introns of the huge gene families fast-evolving *var* genes of *Plasmodium* species). To filter remaining false positives by introns in paralogous genes we first translated all Introner containing genes from DNA to protein sequence. Then we used diamond (version 0.9.24) (*39*), with default options except minimum e-value of 10^-20^, to identify and remove from the list cases of sequence similarity between encoded proteins for intron pairs with similar sequences.

Within each genome with remaining similar intron pairs, we then used pairwise similarities and a greedy algorithm to group introns with at least one remaining pairwise similarity into Introner families. Families with at least 4 introns were retained.

### Filtering assembly errors

We also filtered out putative Introner families identified as a result of genome assembly problems. For example, sequencing adapters included in genome assemblies might sometimes be annotated as introns, in which case genome assemblies including many sequencing adaptors could have multiple similar annotated intronic sequences. We used blast searches to examine putative IntronerIntronerss for the presence of Illumina adaptors and removed families which contained them. We also performed an internet search on each putative Introner family consensus sequence and subsections of each consensus sequence to ensure that Introner families did not embody or contain any known sequences associated with genome assembly or sequencing methods.

### Filtering Introners with low complexity

We filtered putative Introners families with low sequence complexity since sequence similarity between these potential Introners could have resulted from alternative mechanisms than transposition e.g. microsatellite expansion. To do this, we manually examined putative Introner family consensus sequences and looked for an abundance of short repetitive sequence motifs.

### Finalized Introner sequences

After filtering, we possessed a set of finalized Introner sequences sorted in fasta files by species and family within species. Fasta files for each Introner family in each species can be found at https://github.com/lgozasht/Introner-elements.

### A representative set of genomes for downstream analyses based on Introner content

We next scrutinized patterns of Introner family presence/absence across all Introner-containing genomes. In particular, the presence of clusters of closely-related organisms represented within the TARA Oceans genomes led to cases of multiple genomes with very similar Introner complements -- both at the level of families and of specific insertions. At the same time, as mentioned in the main text, closely-related genomes sometimes exhibit overlapping but nonidentical sets of Introner families. Genomes containing identical sets of Introner families were grouped, leading to 16 groups, mostly including 2 genomes (13 groups), but ranging up to 9 genomes.

### Proportion of Introners inside of genes

Since Introners in intergenic regions cannot be annotated as introns, we developed a systematic method to re-cover them conditional on a known Introner family detected as described above. We employed multiple alignment using fast Fourier transform (MAFFT) (*40*) to conduct multiple sequence alignments for each Introner family in each species. Next, we generated a consensus sequence for each Introner family using a positional nucleotide frequency matrix. We required that greater than 50% of Introners possess the same nucleotide at a particular position for that base to be included in our consensus sequence. We BLASTed each consensus to its corresponding reference genome and filtered duplicate and self hits.

We used a permutation test to interrogate possible enrichment for Introners in genes. If a genome is more gene dense, a transposon is more likely to land in a gene. To correct for this, we generated 1000 permutations for the probability that a particular Introner will insert into a gene by chance by randomizing the Introner positions across the genome such that n = the number of total insertions and p = gene density. We compared these with our actual values to test for insertional enrichment in genic regions.

### GC content analysis

Genes are generally GC rich relative to intergenic regions (*41*), and transposable elements have been shown to display preference for GC rich regions (*42, 43*). To test whether Introners that are enriched in genes also demonstrate a bias for GC rich regions, we employed a permutation test. We calculated the GC content for the concatenated 10 base pairs (bp) upstream and downstream of each insertion (20bp total). We then generated 10000 permutations for the GC content of randomly resampled 20 bp regions from the same gene in which each respective Introner was found. We compared our observed GC proportions to the randomly sampled distribution and found that Introners in many species are also enriched within GC rich regions. Across all species, we found a significant correlation between insertional enrichment in genes and insertional enrichment in GC rich regions, suggesting that Introners may favor GC rich regions rather than simply an insertion preference for genic regions *per se* (P < 0.0001; binomial test). Here, we constructed this comparison as a one-sided test where we asked if the proportion of species whose insertion preferences exceeded background genic GC content differed from random expectations. Note also that conditioning on the specific genes into which Introners insert is very conservative because any strong GC content skew within a gene would be reflected in the null distribution.

### Nucleosome occupancy prediction and analysis

Since the vast majority of genomes in our survey lack available epigenetic data and most cannot be reliably cultured, we applied an *in silico* predictive approach to interrogate whether Introners insert into nucleosome linker DNA in other lineages. We used the program, *NuPoP* (*44*), to predict the nucleosome occupancy for the 10kb spanning and surrounding the insertion sites of all Introners in each species for which we possessed at least 4 Introners with contiguous sequence assembled 5kb upstream and 5kb downstream of insertion sites. We required at least 5kb around each Introner to ensure that we maximize the accuracy of predictions. The reason is that this approach is based on a hidden Markov model and therefore requires moderate sequence lengths to produce reliable results. We ran *NuPoP* with flags *species=0* and *model=4* first with Introners and then again with Introners computationally removed from the gene sequence as a control (Supplementary Figure 2). We observe a pattern across most genomes of decreased nucleosome occupancy for the 100 bp surrounding the 5’ splice sites of Introners relative to background regions (p = 8.26e-07; binomial test), and are able to reproduce the pattern previously reported for Introners in Micromonas and Aerococcus using nucleosome profile data (Supplementary Figure 2-3). We observe a decreased nucleosome occupancy within Introner sequences relative to background regions even more often (p = 4.00e-10; binomial test), suggesting that Introners insert into and inhabit nucleosome linker regions. When we perform the same comparison with Introners removed to replicate surrounding regions, we observe the opposite pattern, in which the nucleosome occupancy is higher (p = 9.13e-12; binomial test), suggesting that nucleosomes often flank Introner sequences (Supplementary Figure 3). We used a gaussian GLM through R to look for an association between the number of Introners and predicted nucleosome occupancy within Introners relative to background nucleosome occupancy of each Introner family in each species with formula *delta_nuc_occup ~ number_of_Introners*. We did the same association for Introner length and species with formulae: *delta_nuc_occup ~ Introner_length* and *delta_nuc_occup ~ species.* We find that the number of Introners in each family and Introner length are both good predictors of nucleosome occupancy within Introners relative to background (P < 0.0001 and P = 0.04), with smaller families with shorter sequences having lower *delta_nuc_occup*. We postulate that this association may be due to reduced accuracy of predictions on small sample sizes. This may explain why in smaller, shorter Introner families we sometimes don’t observe as clear of a pattern of low nucleosome occupancy in Introners. We note however, that species is an even better predictor than the aforementioned variables (P < 0.0001), and that our ability to predict nucleosome profiles accurately with a sequence motif based HMM could also be limited in highly divergent species relative to those used for training the HMM initially or in low quality genomes. Indeed, when we filter for species in which we observed *delta_nuc_occup* < 0, and use a GLM with the formula: *delta_nuc_occup ~ GC_content,* we find that GC content explains better than species (P = 0.0015).

### Identifying TSDs and TIRs

We searched for evidence of TSDs in and around each Introner family in each species for which there were at least 15 genic Introners in the family. Most clearly, positions that are part of a TSD manifest as similarity between 5’ and 3’ ends within individual Introner from a single insertion site, but not globally between introns (that is, corresponding nucleotides 5’ and 3’ ends match for intron A, but do not necessarily match between introns A and B). However, TSDs often include core GY/AG splice site nucleotides, in which case nucleotides will match across introns as well. Thus to assess TSD presence/absence we calculated both absolute match between corresponding 5’ and 3’ nucleotides (that is, nucleotides the same distance from the splice site, for instance the first nucleotide of the intron and the first nucleotide of the downstream exon), as well as relative match, calculated as the fraction of matches within individual introns divided by the expected value calculated from 1000 random 5’/3’ pairs. For each Introner family within each species, these values were assessed to look for TSD patterns including at least one nucleotide position with locus-specific match (e.g., NAGgy...nag, where N/n show significant match within but not between insertion sites). For patterns that included only across-intron matches (e.g., families where all or nearly all introns are preceded by an AG (AGgy...xyag), it is not possible to distinguish between the AG representing a TSD (Introner sequence = gy..xyag, TSD=AG), or the AG representing an insertion site without a TSD (Introner sequence = gy..xyag or AGgy...xy, no TSD). In some instances, one of the two duplicate motifs was part of an extended TIR (see below). Otherwise, TSD presence/absence was called as ambiguous.

TIRs were searched for manually by searching the consensus sequence within a region extending from −20 to 20 nucleotides of the intron. This generally yielded either a clear extended TIR (≥6 nucleotides) or no evidence of a TIR. The few cases with partial or short TIRs were called as ambiguous. These calls are available at https://github.com/lgozasht/Introner-elements

### Examples used in Figure 2A-F

For Figure 2A, we used an Introner in family 1 of *Alternaria alternata* (Genbank acc. GCA_001572055.1) on scaffold LPVP01000001.1, position 1639687-1639750. For Figure 2B, we used an Introner in family 1 of *Symbiodinium microadriaticum* (Genbank acc. GCA_001939145.1) on scaffold LSRX01000224.1 position 86725-86839. For Figure 2C, we used an Introner in family 5 of *Chrysochromulina sp.* (TARA_PON_109_MAG_00232) on scaffold 000000000114 position 1755-1824. For Figure 2D, we used an Introner in family 4 of *Acanthoeca sp.* (TARA_AON_82_MAG_00310) on scaffold 000000000270.1.1.3 at position 451-559. For Figure 2E, we used an Introner in family 2 of *Florenciella sp.*

(TARA_MED_95_MAG_00409) on scaffold 000000000986.1.2.2 at position 852-918. For figure 2F, we used an Introner in family 4 of *Aureococcus sp.* (TARA_AOS_82_MAG_00129) on scaffold 000000001056 at position 3289-3365.

### Comparing orthologs between aureococcus isolates

We used BLAST to identify homology between Introner containing genes in *Aureococcus sp.* (isolate TARA_AOS_82_MAG_00129) and *Aureococcus anophagefferens* (Genbank Acc. GCA_000186865.1). We used MAFFT (*40*) to perform a multiple sequence alignment (MSA) between each Introner containing gene in *Aureococcus sp.* and its match in *Aureococcus anophagefferens* with the lowest e-value given the e-value < 0.01. We also performed an msa between translated proteins corresponding to these genes. The example shown in Figure 2 stems from an alignment between an Introner containing gene in *Aureococcus sp.* and *Aureococcus anophagefferens* Phosphoenolpyruvate carboxylase kinase 1 (NCBI Acc. XM_009038642.1) at both the nucleotide and protein level. In this example, *Aureococcus sp.* exhibits an Introner insertion at position 545 in this gene, which resulted in the addition of 4 amino acids relative to Aureococcus anophagefferens.

### Identifying potential mobilizing elements

We searched for transposase-encoding autonomous elements that may mobilize Introners with similar terminal sequences. For each Introner family in each species, we constructed position weight matrices of length 22 bp at the 5’ and 3’ ends of the elements. We then searched each respective species’ genome for matches to the 5’ PWM using PoSSuMsearch (*45*) and searched the downstream 10,000 bp of each match using the 3’ PWM. We also searched for matches among predicted repetitive elements found using RepeatModeler (*46*). Open reading frames were found between the 5’-3’ pairs of PWM matches and their translated amino acid sequences were used to search a database of transposases (a subset of UniProt) using BLASTP (Supplementary Table 8).

### Assessing homology between Introners in different species

We used BLAST to search for homology between consensus sequences of Introners in different species. We performed an all vs all BLAST of Introner consensi and found no evidence of homology except between relatively recently diverged species (within the same genus). However, we did observe cases for TARA MET genomes (for which only the genus is reported) in which isolates within the same genus possess different Introner families. We treated those isolates as separate species throughout our study.

### Checking for associations between TSD and TIR presence and Introner architecture and distribution

We tested for associations between TSD and TIR presence and other Introner statistics both on the subset of genomes that possessed Introner families with and without TSDs/TIRs and across all species. To do this, we used generalized linear models through R (Supplementary Table 7). To test for an association between TSD and TIR presence and insertional preference in genes, we fit a GLM with the following format:

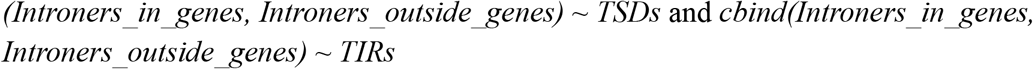

using a binomial family link function. As a control, performed the same analysis with species instead of TSD or TIR presence using a GLM of the format:

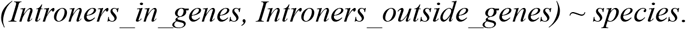

We found that the term species better explains our data than TSD or TIR presence using AIC.

To test for an association between TSDs and TIR and canonical splice site usage, we fit a GLM of the form:

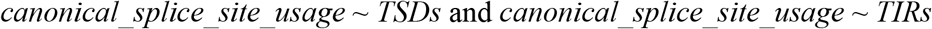

under the gaussian family link function. Again we performed the same association for species and found that species better explains our data than TSD or TIR presence. To test for an association between TSD and TIR presence and number of Introners, we used a GLM of the form:

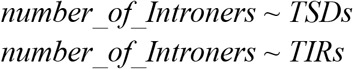

under a gaussian link function. We find that TSDs and TIRs explain our data poorly in this case. To test for an association between TSD and TIR presence and delta nucleosome occupancy, we a GLM of the format:

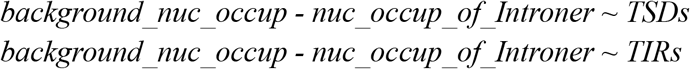

again using the gaussian family link function. We performed the same association with species and again found that species better explained our data.

### RNA-seq analysis

For each species with identified Introners (genus level TARA metagenomes excluded), we searched the SRA database for RNA-seq data, prioritizing sequencing runs conducted on the same individual from which the reference genome was assembled (Supplementary Table 9). We aligned the RNA reads to the reference genome using *STAR* (*47*), calculated the depth at each site using *samtools* (*48*), and identified splice junctions using *leafcutter* (*49*). We then used custom python scripts to identify, for each intron, the number of splicing events that used the annotated splice junctions, as well as the number of splicing events that used non-canonical junctions within 50 nucleotides on either side of the annotated junction. We then used the R package *lme4* (*50*) to construct generalized linear model of the form:

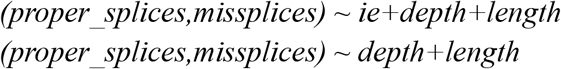

to correct for the depth and length of each intron. To ensure that the *ie* variable (whether or not the intron is an Introner) was significantly correlated with splicing behavior, we calculated the likelihood ratio of the two models using the Akaike Information Criterion (AIC) (*51*). If the model containing the *ie* variable was a better fit, and if coefficient for the *ie* variable was positive, Introners in this species exhibit more canonical splicing than non-Introner intron.

We used a similar approach for percent spliced in (PSI; Fig. 3), using GLMs of the form:

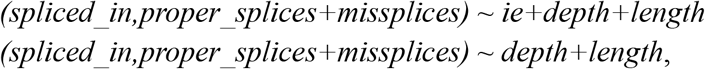

ensuring that whether an intron is an Introner is significantly correlated with PSI using AIC likelihood ratios, and considering cases with negative coefficients for the *ie* variable as cases where Introners are more likely to be spliced in than non-Introner introns (Fig. 3).

### Correlating aquatic lifestyle and Introner presence

For phylogenetic tests, we downloaded the global tree from *open tree of life* (*52*) and pruned the tree to retain only species that we considered in this analysis. In the case of TARA MET genomes, for which we only possessed the genera, we randomly selected one species within each genus to represent all isolates. Our tree can be found at https://github.com/lgozasht/Introner-elements/blob/main/pruned_tree2.nwk.gz. To evaluate a correlation between aquatic lifestyle and the presence of Introners, we used both Pagel’s test and a binary phylogenetic generalized mixed model in the R package *phytools* (*53, 54*) using the aforementioned global phylogeny as an input (Supplementary Figure 5). Pagel’s test constructs models in which the two traits evolve independently, co-dependently, or with one trait dependent upon the other, and model fits are compared using AIC (Supplementary Figure 4). The binary phylogenetic generalized mixed model performs a linear regression, estimates the strength of phylogenetic signal, and conducts an approximate conditional likelihood test. To prepare the input data for these tests, we manually annotated each species considered as aquatic or not using the following criterion: To be considered an aquatic species, a species must spend most of its life surrounded by water (Supplementary Table 4). A species also cannot be an obligate parasite of a terrestrial organism even though in such a case the species may live primarily within an aqueous solution.

