## Supplementary Figures for "Transposable elements drive intron gain in diverse eukaryotes"

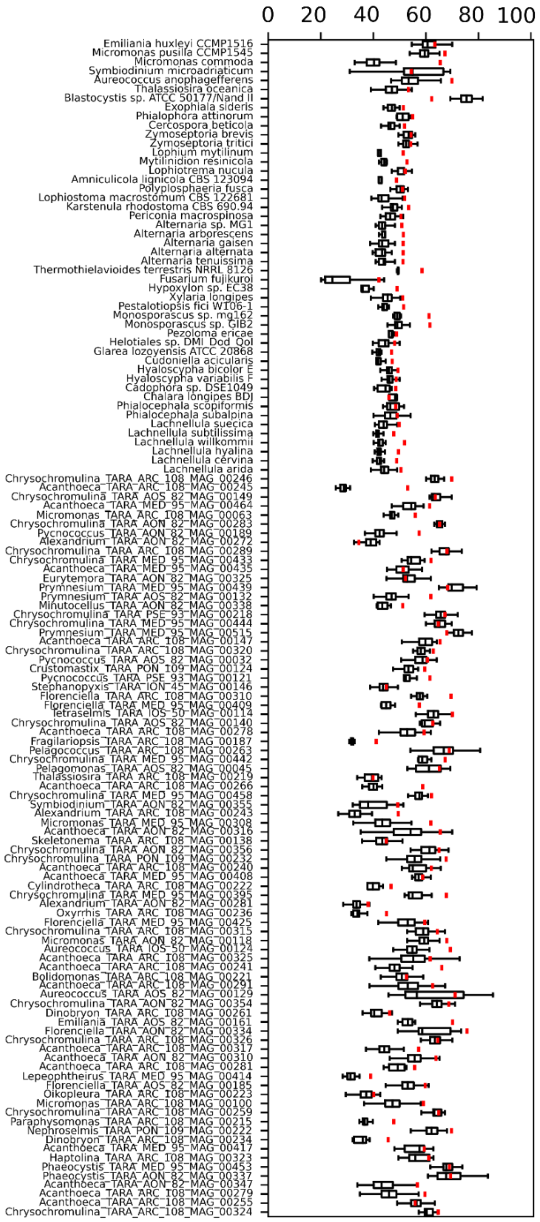


**Supplementary Figure S1: Insertion of Introners into GC rich regions.** Box plots represent 1000 permutations for the GC content of possible insertion sites within Introner containing genes as determined by randomization. Red bars denote GC content of the 10bp surrounding observed insertion sites.

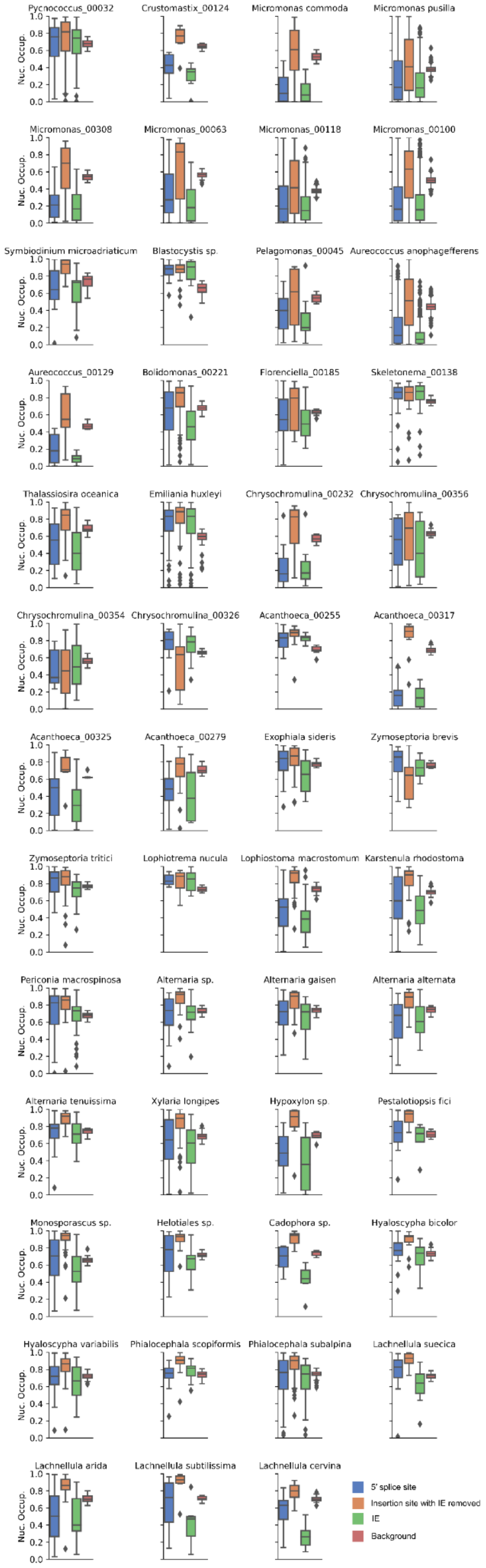

Pycnococcus_00032
Crustomastix_00124
Micromonas commoda
Micromonas pusilla
Micromonas_00308
Micromonas_00063
Micromonas_00118
Micromonas_00100
Symbiodinium microadriaticum
Blastocystis sp.
Pelagomonas_00045
Aureococcus anophagefferens
Aureococcus_00129
Bolidomonas_00221
Florenciella_00185
Skeletonema_00138
Thalassiosira oceanica
Emiliania huxleyi
Chrysochromulina_00232
Chrysochromulina_00356
Chrysochromulina_00354
Chrysochromulina_00326
Acanthoeca_00255
Acanthoeca_00317
Acanthoeca_00325
Acanthoeca_00279
Exophiala sideris
Zymoseptoria brevis
Zymoseptoria tritici
Lophiotrema nucula
Lophiostoma macrostomum
Karstenula rhodostoma
Periconia macrospinosa
Alternaria sp.
Alternaria gaisen
Alternaria alternata
Alternaria tenuissima
Xylaria longipes
Hypoxylon sp.
Pestalotiopsis fici
Monosporascus sp.
Helotiales sp.
Cadophora sp.
Hyaloscypha bicolor
Hyaloscypha variabilis
Phialocephala scopiformis
Phialocephala subalpina
Lachnellula suecica
Lachnellula arida
Lachnellula subtilissima
Lachnellula cervina
Nuc. Occup.
0.0
0.2
0.4
0.6
0.8
1.0
5' splice site
Insertion site with IE removed
IE
Background

**Supplementary Figure 2: Nucleosome occupancy with Introners present and Introners removed**.

Box plots represent predicted nucleosome occupancy distributions for different regions related to Introners for all Introners in each species. Colors denote the region, with blue showing the nucleosome occupancy for the 100bp surrounding the 5' splice site, orange showing nucleosome occupancy for the 100bp surrounding the Introner with the Introner removed, green showing the nucleosome occupancy across Introner sequences, and red showing the background nucleosome occupancy. In most species, we find that the 100bp surrounding the 5' splice site of Introners exhibits lower nucleosome occupancy relative to background regions (p = 8.26e-07; binomial ). We observe the same pattern for Introner sequences (p = 4.00e-10; binomial). When we remove Introners, we instead observe the opposite pattern, in which nucleosome occupancy is high relative to the background, suggesting that Introners nucleosomes are often bordered by nucleosomes (p = 9.13e-12; binomial).

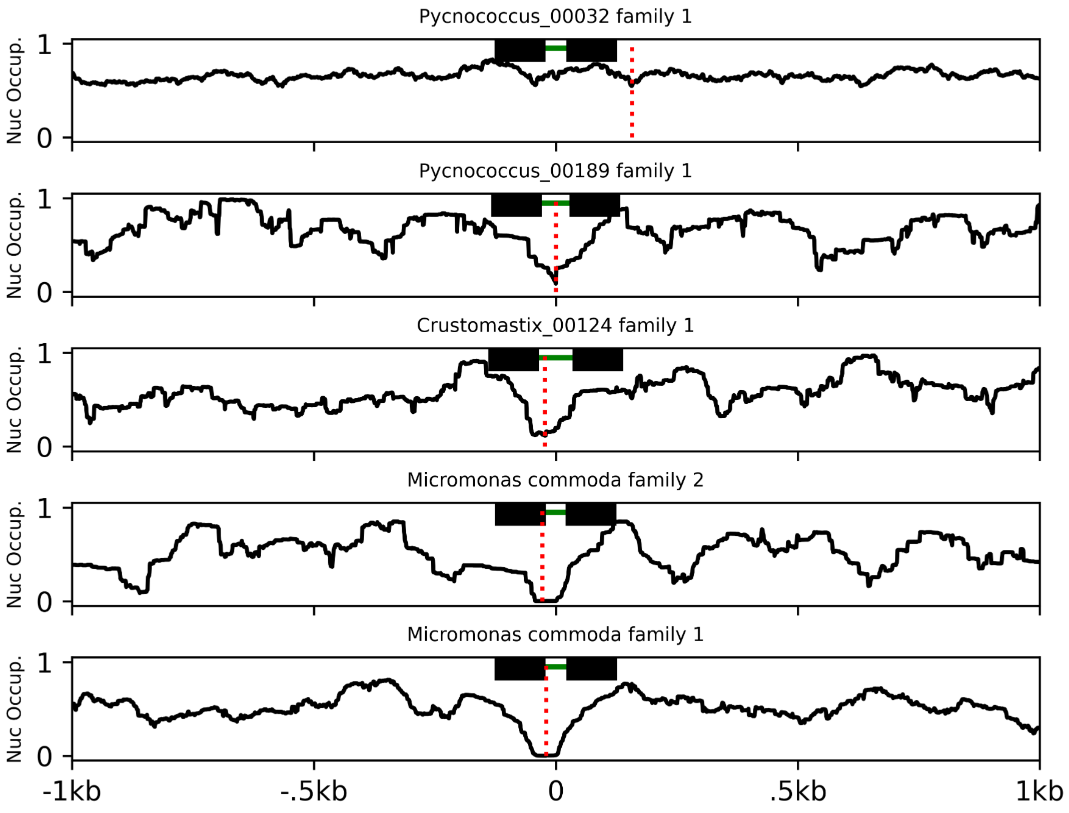

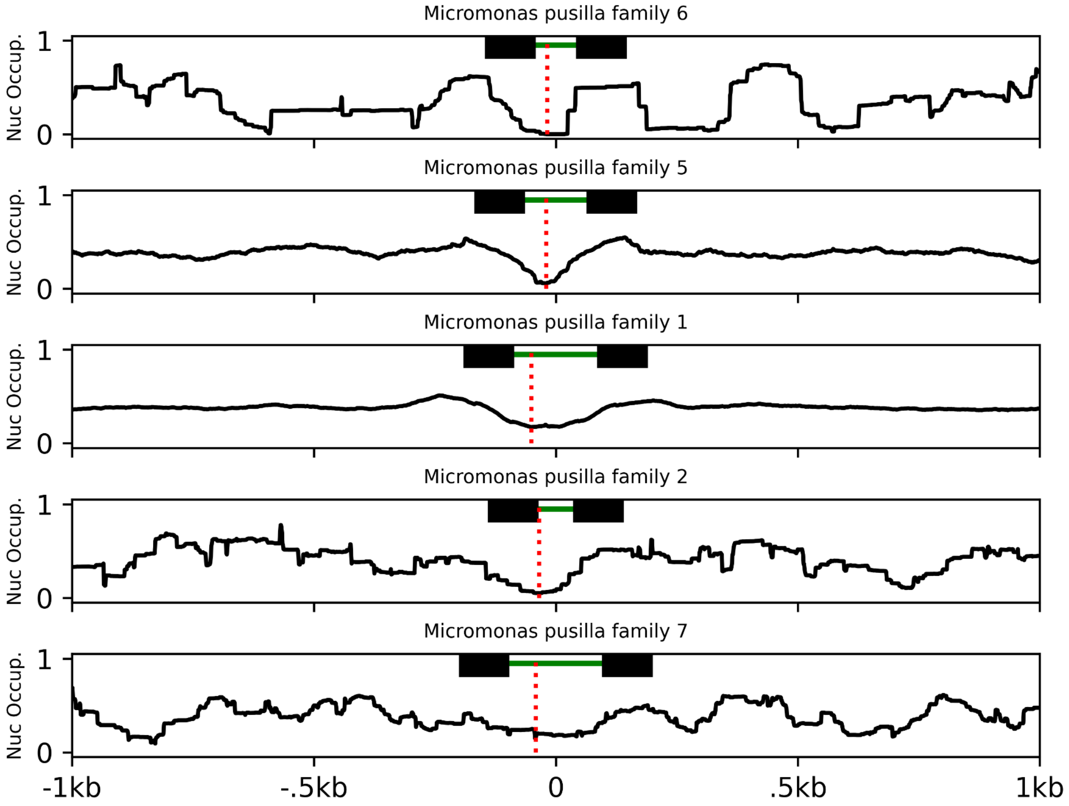
Micromonas pusilla family 6
Micromonas pusilla family 5
Micromonas pusilla family 1
Micromonas pusilla family 2
Micromonas pusilla family 7
Nuc Occup.
1
0
-1kb
-.5kb
0
.5kb
1kb

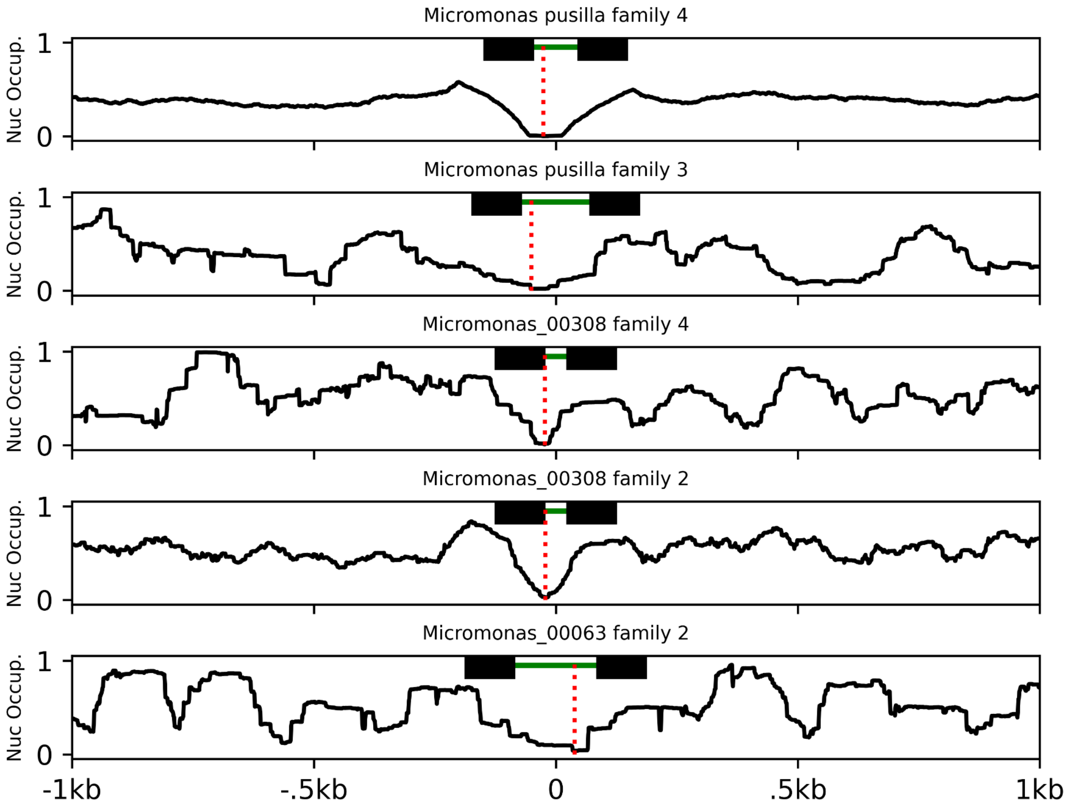

Micromonas pusilla family 4
Micromonas pusilla family 3
Micromonas_00308 family 4
Micromonas_00308 family 2
Micromonas_00063 family 2
Nuc Occup.
1
0
-1kb
-.5kb
0
.5kb
1kb

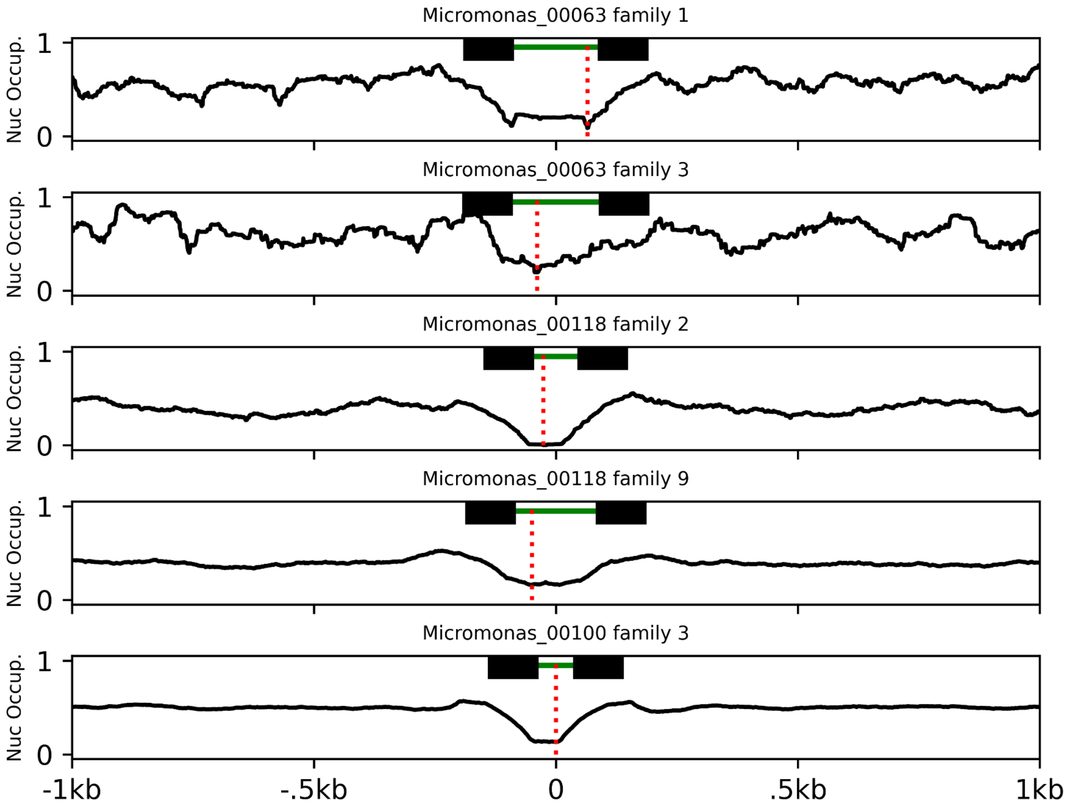

Micromonas_00063 family 1
Micromonas_00063 family 3
Micromonas_00118 family 2
Micromonas_00118 family 9
Micromonas_00100 family 3
Nuc Occup.
1
0
-1kb
-.5kb
0
.5kb
1kb

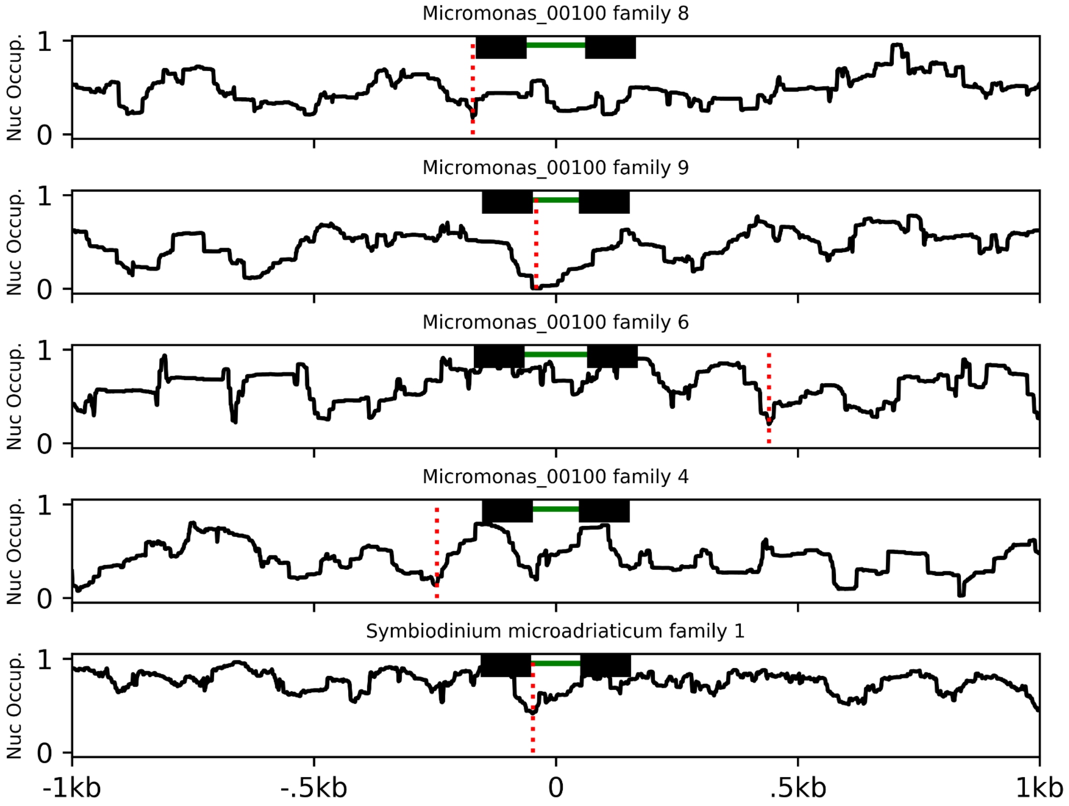
Micromonas_00100 family 8
Micromonas_00100 family 9
Micromonas_00100 family 6
Micromonas_00100 family 4
Symbiodinium microadriaticum family 1
Nuc Occup.
1
0
-1kb
-.5kb
0
.5kb
1kb

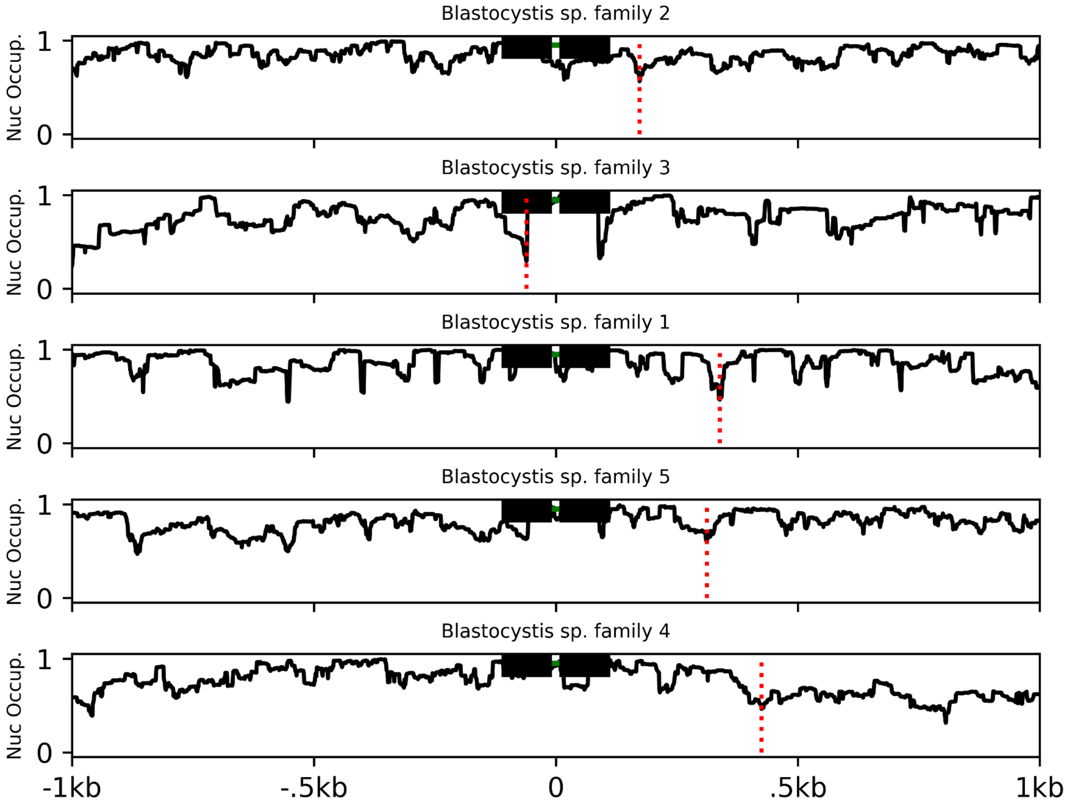

Blastocystis sp. family 2
Blastocystis sp. family 3
Blastocystis sp. family 1
Blastocystis sp. family 5
Blastocystis sp. family 4
Nuc Occup.
1
0
-1kb
-.5kb
0
.5kb
1kb

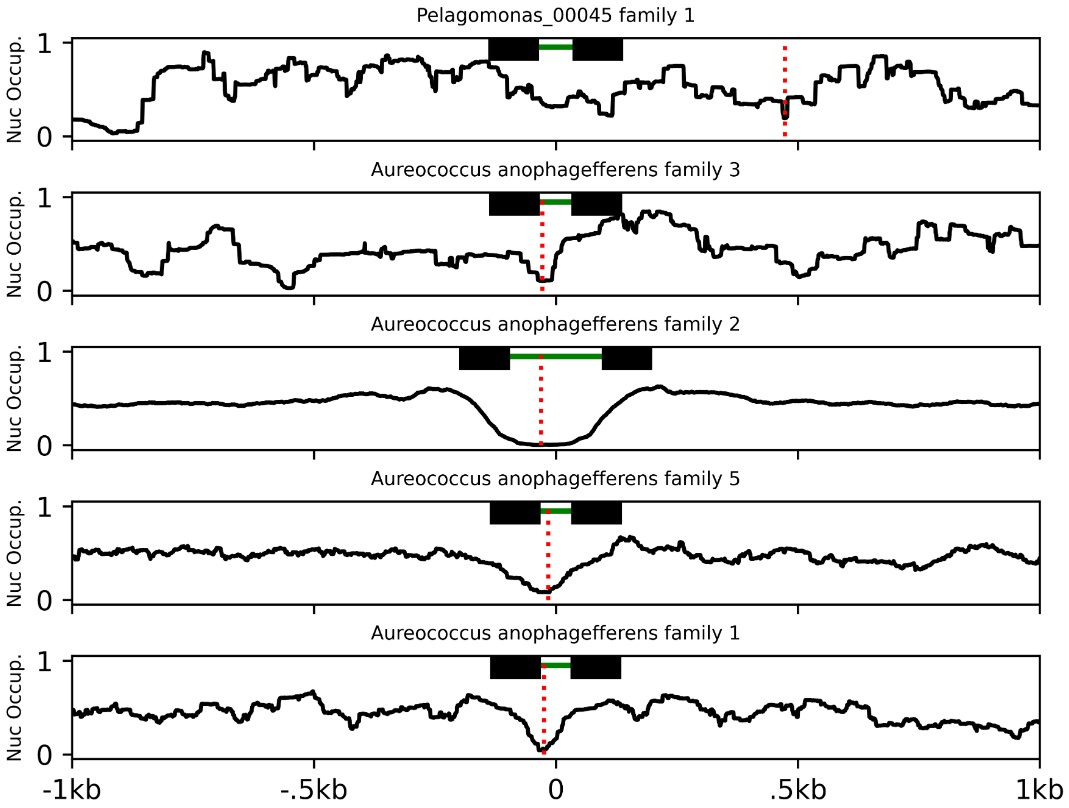

Pelagomonas_00045 family 1
Aureococcus anophagefferens family 3
Aureococcus anophagefferens family 2
Aureococcus anophagefferens family 5
Aureococcus anophagefferens family 1
Nuc Occup.
1
0
-1kb
-.5kb
0
.5kb
1kb

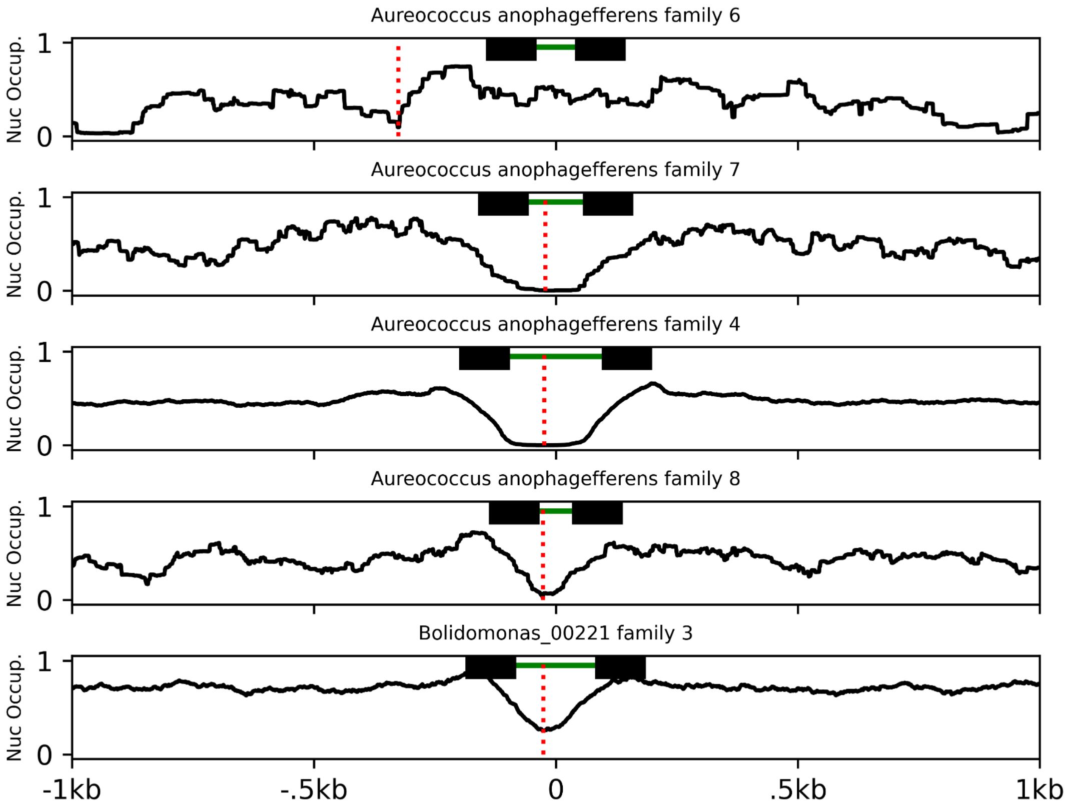

Aureococcus anophagefferens family 6
Aureococcus anophagefferens family 7
Aureococcus anophagefferens family 4
Aureococcus anophagefferens family 8
Bolidomonas_00221 family 3
Nuc Occup.
1
0
-1kb
-.5kb
0
.5kb
1kb

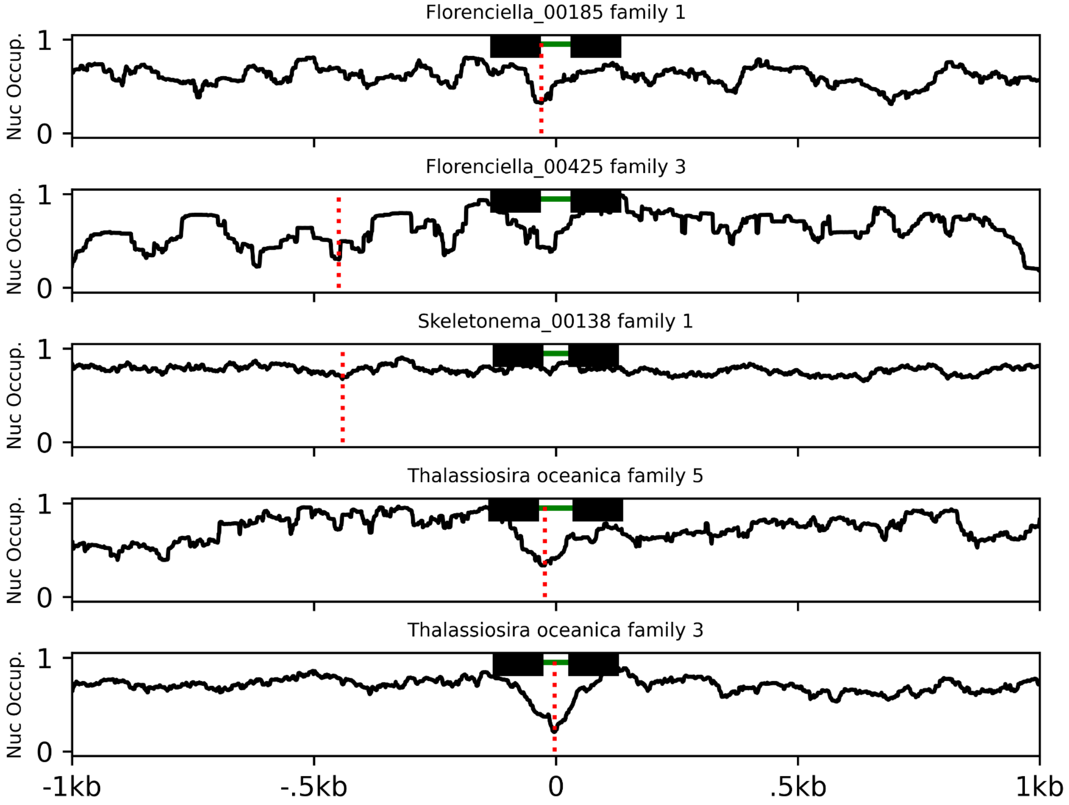

Florenciella_00185 family 1
Florenciella_00425 family 3
Skeletonema_00138 family 1
Thalassiosira oceanica family 5
Thalassiosira oceanica family 3
Nuc Occup.
1
0
-1kb
-.5kb
0
.5kb
1kb

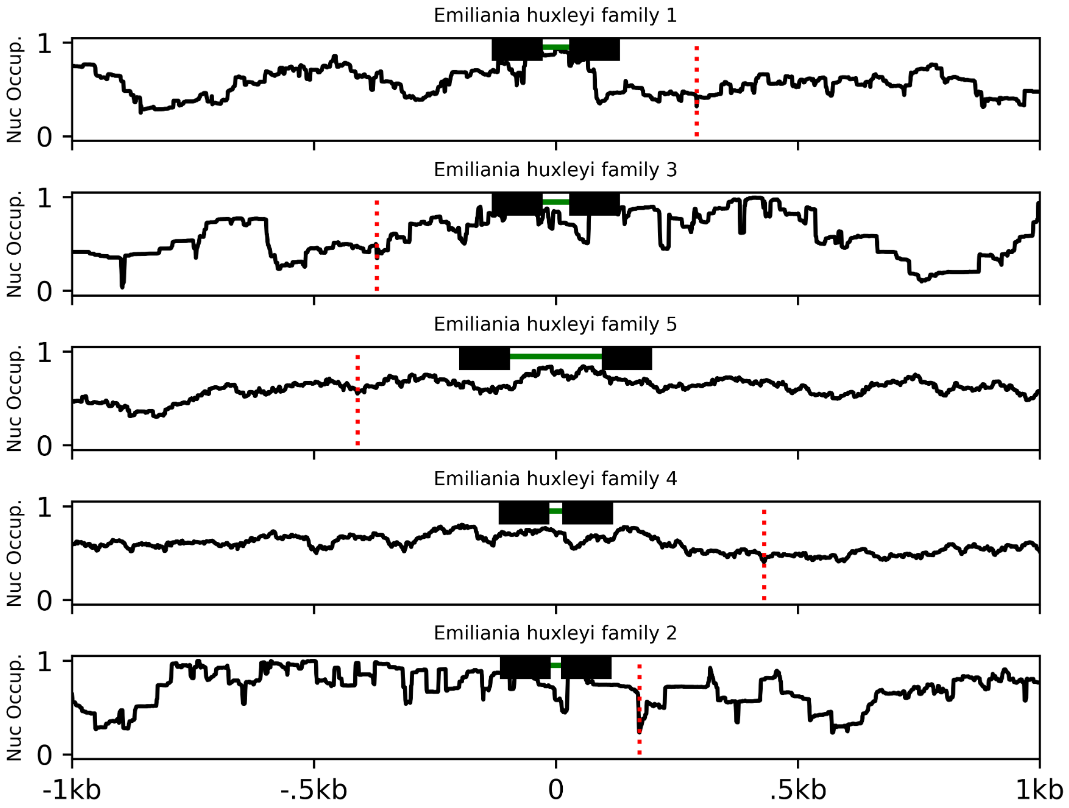

Emiliania huxleyi family 1
Emiliania huxleyi family 3
Emiliania huxleyi family 5
Emiliania huxleyi family 4
Emiliania huxleyi family 2
Nuc Occup.
1
0
-1kb
-.5kb
0
.5kb
1kb

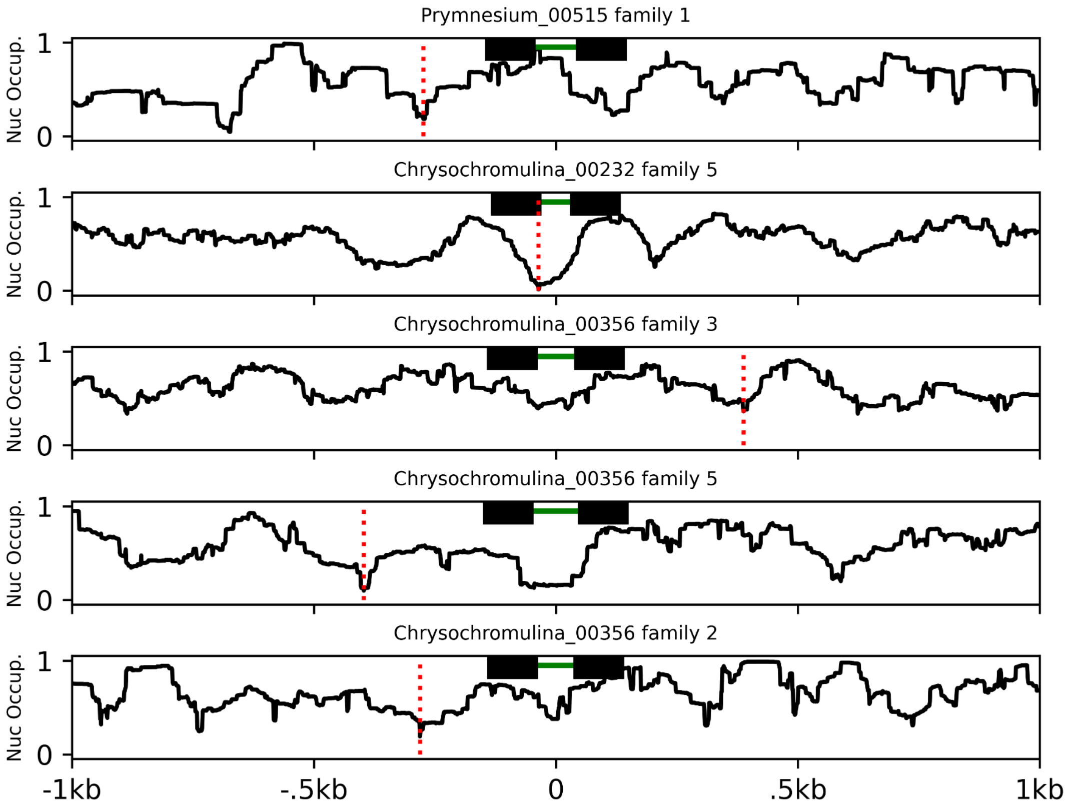
Prymnesium_00515 family 1
Chrysochromulina_00232 family 5
Chrysochromulina_00356 family 3
Chrysochromulina_00356 family 5
Chrysochromulina_00356 family 2
Nuc Occup.
1
0
-1kb
-.5kb
0
.5kb
1kb

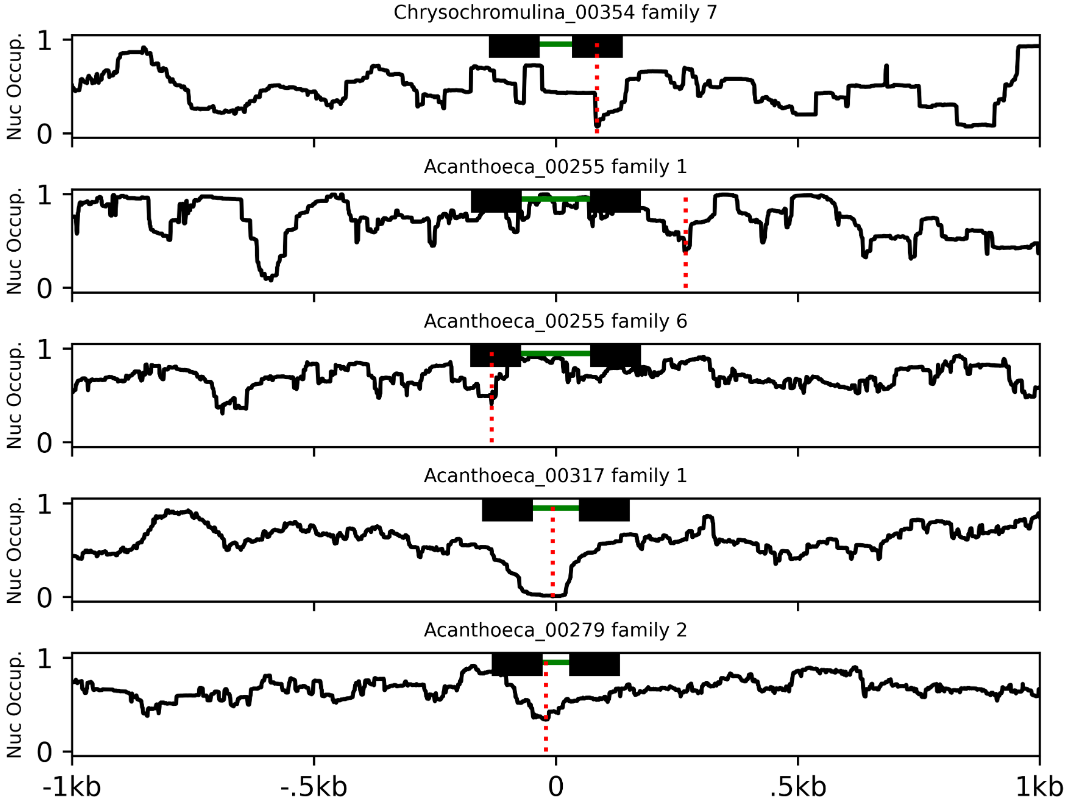
Chrysochromulina_00354 family 7
Acanthoeca_00255 family 1
Acanthoeca_00255 family 6
Acanthoeca_00317 family 1
Acanthoeca_00279 family 2
Nuc Occup.
1
0
-1kb
-.5kb
0
.5kb
1kb

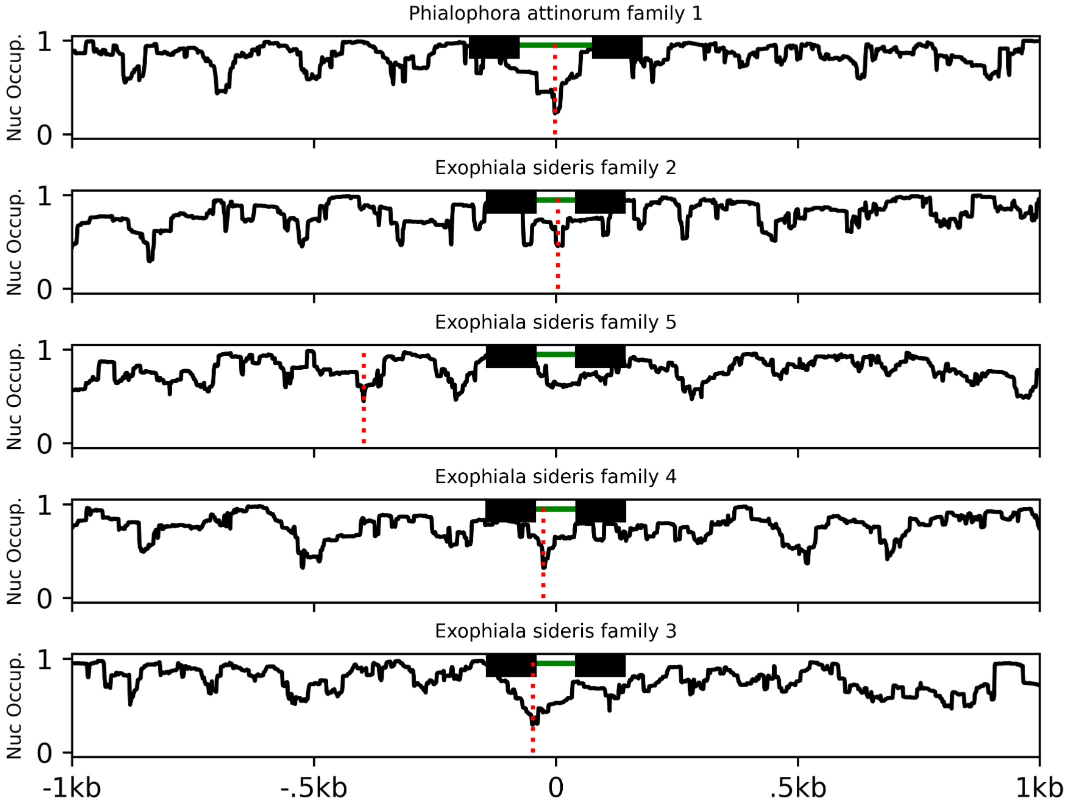

Phialophora attinorum family 1
Exophiala sideris family 2
Exophiala sideris family 5
Exophiala sideris family 4
Exophiala sideris family 3
Nuc Occup.
1
0
-1kb
-.5kb
0
.5kb
1kb

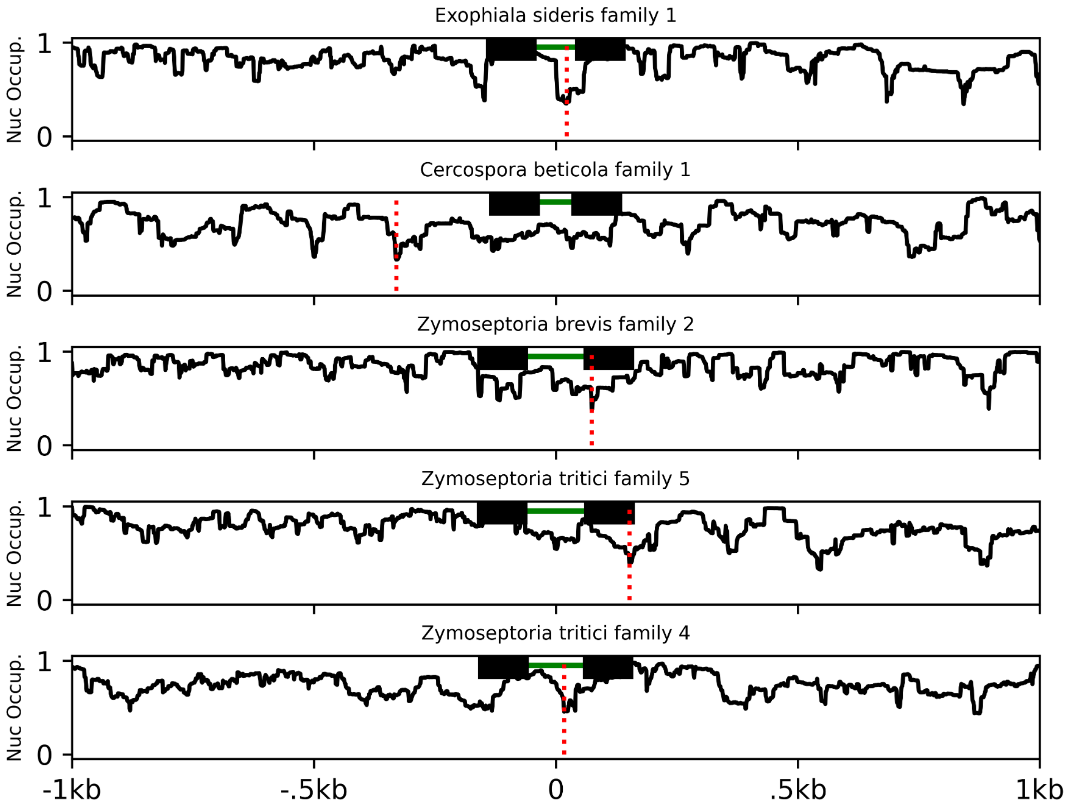
Exophiala sideris family 1
Cercospora beticola family 1
Zymoseptoria brevis family 2
Zymoseptoria tritici family 5
Zymoseptoria tritici family 4
Nuc Occup.
1
0
-1kb
-.5kb
0
.5kb
1kb

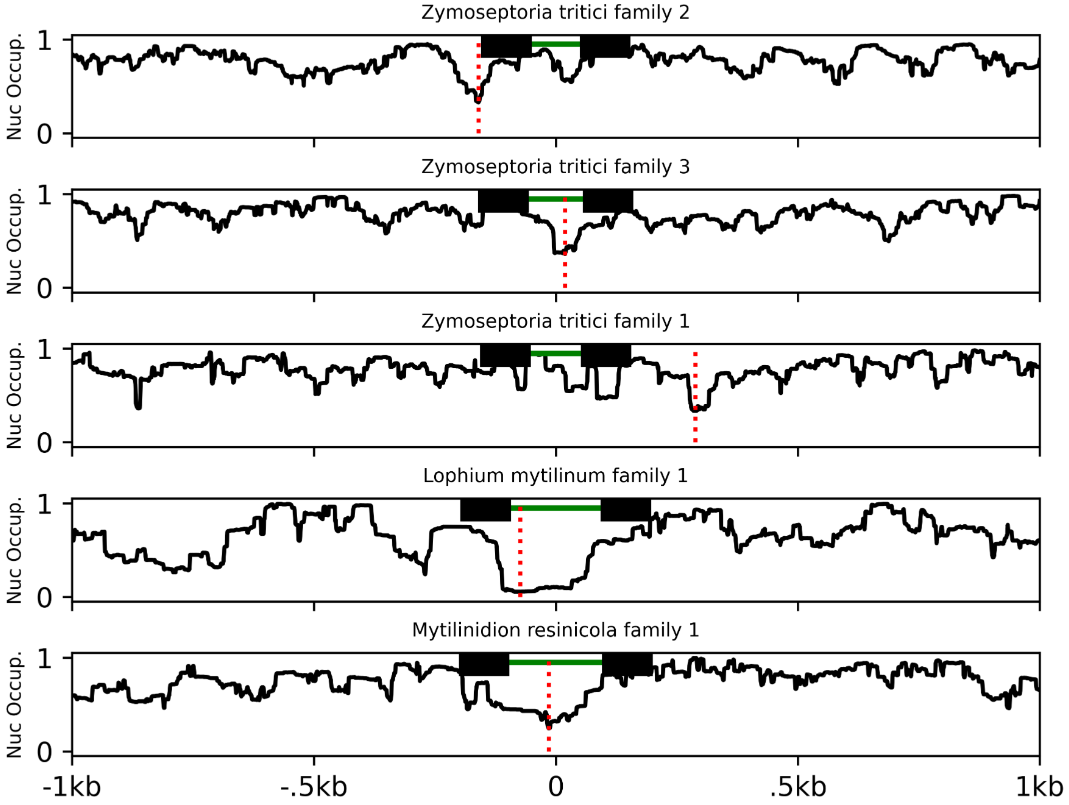

Zymoseptoria tritici family 2
Zymoseptoria tritici family 3
Zymoseptoria tritici family 1
Lophium mytilinum family 1
Mytilinidion resinicola family 1
Nuc Occup.
1
0
-1kb
-.5kb
0
.5kb
1kb

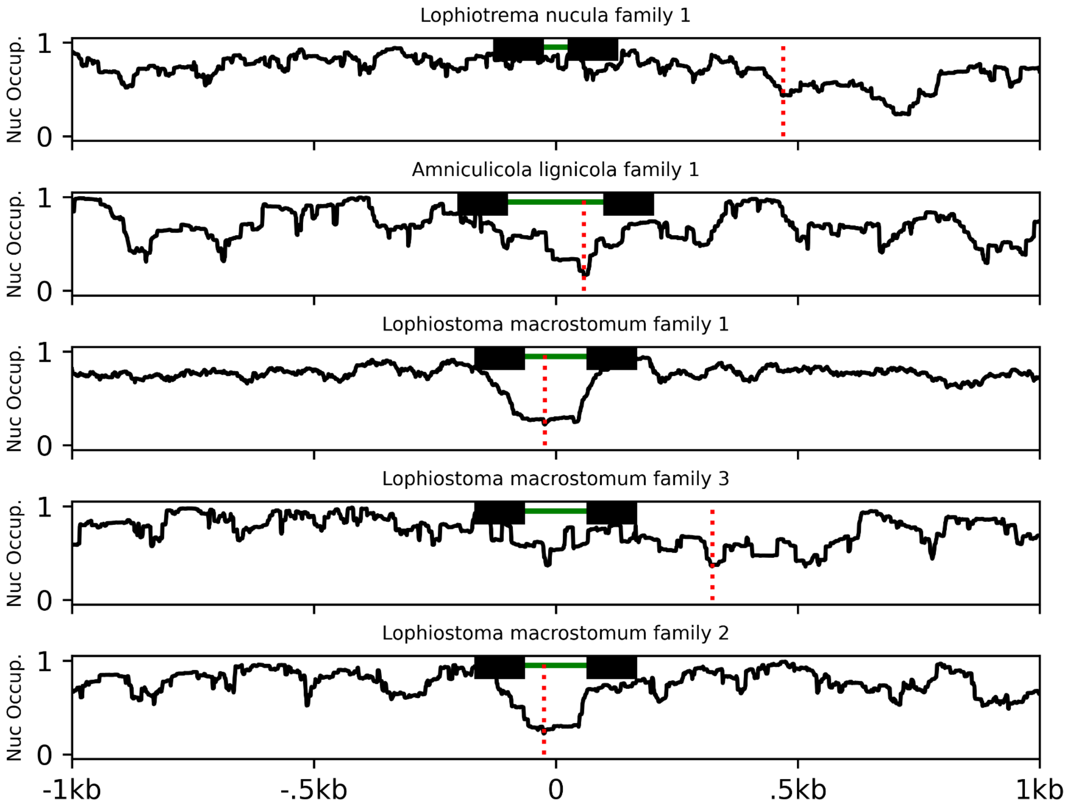

Lophiotrema nucula family 1
Amniculicola lignicola family 1
Lophiostoma macrostomum family 1
Lophiostoma macrostomum family 3
Lophiostoma macrostomum family 2
Nuc Occup.
1
0
-1kb
-.5kb
0
.5kb
1kb

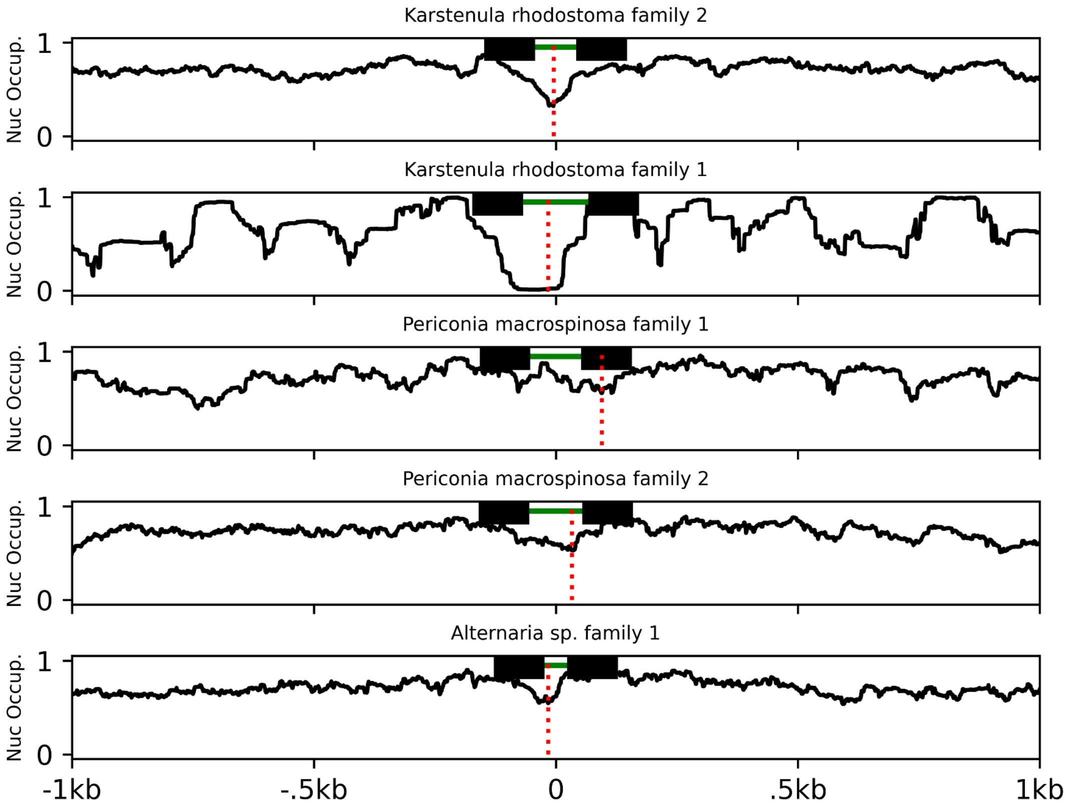

Karstenula rhodostoma family 2
Karstenula rhodostoma family 1
Periconia macrospinosa family 1
Periconia macrospinosa family 2
Alternaria sp. family 1
Nuc Occup.
1
0
-1kb
-.5kb
0
.5kb
1kb

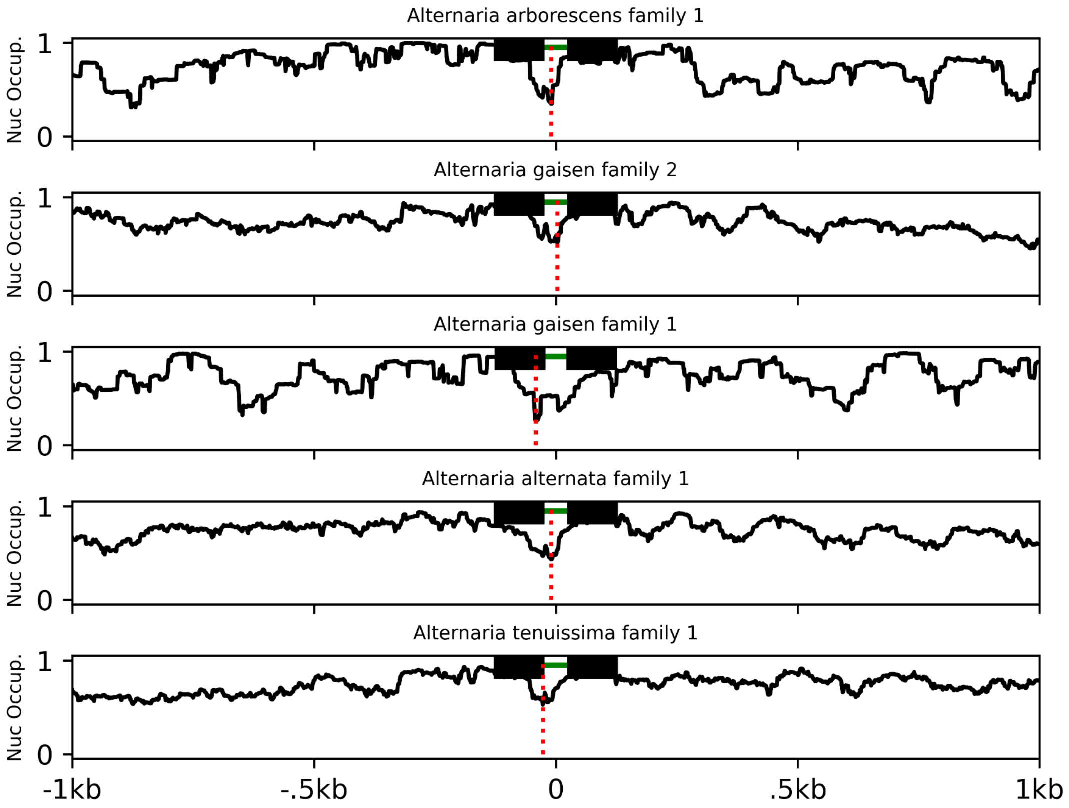
Alternaria arborescens family 1
Alternaria gaisen family 2
Alternaria gaisen family 1
Alternaria alternata family 1
Alternaria tenuissima family 1
Nuc Occup.
1
0
-1kb
-.5kb
0
.5kb
1kb

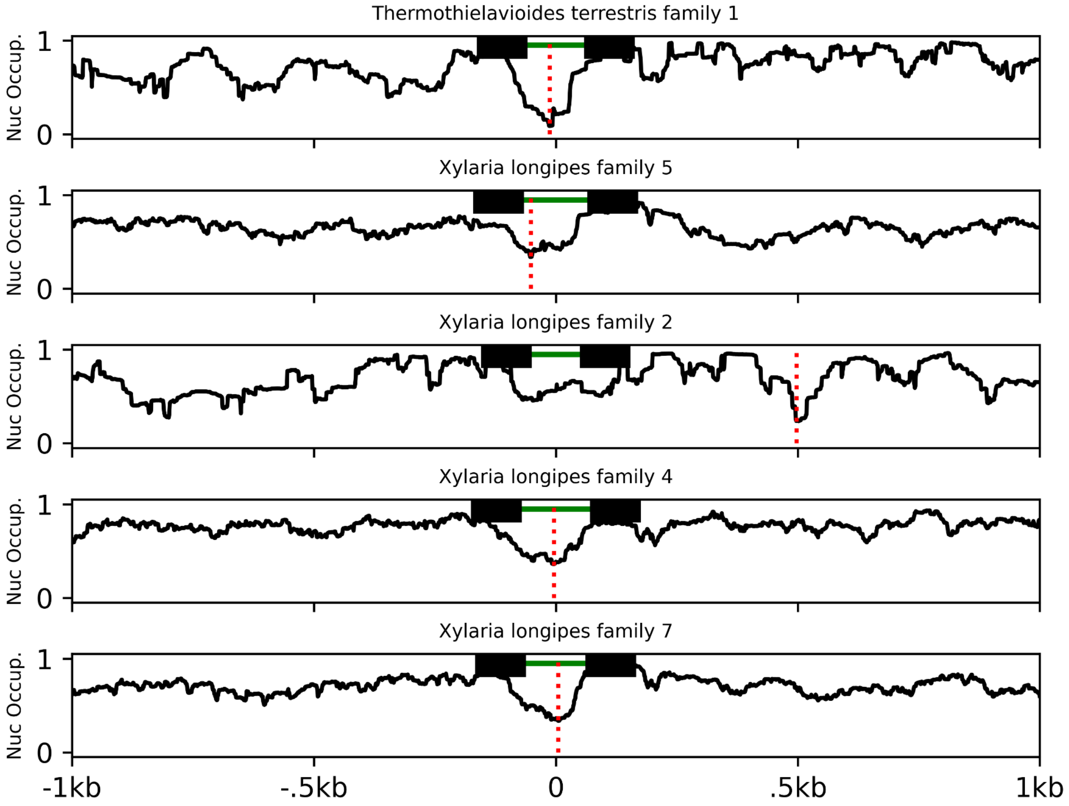
Thermothielavioides terrestris family 1
Xylaria longipes family 5
Xylaria longipes family 2
Xylaria longipes family 4
Xylaria longipes family 7
Nuc Occup.
1
0
-1kb
-.5kb
0
.5kb
1kb

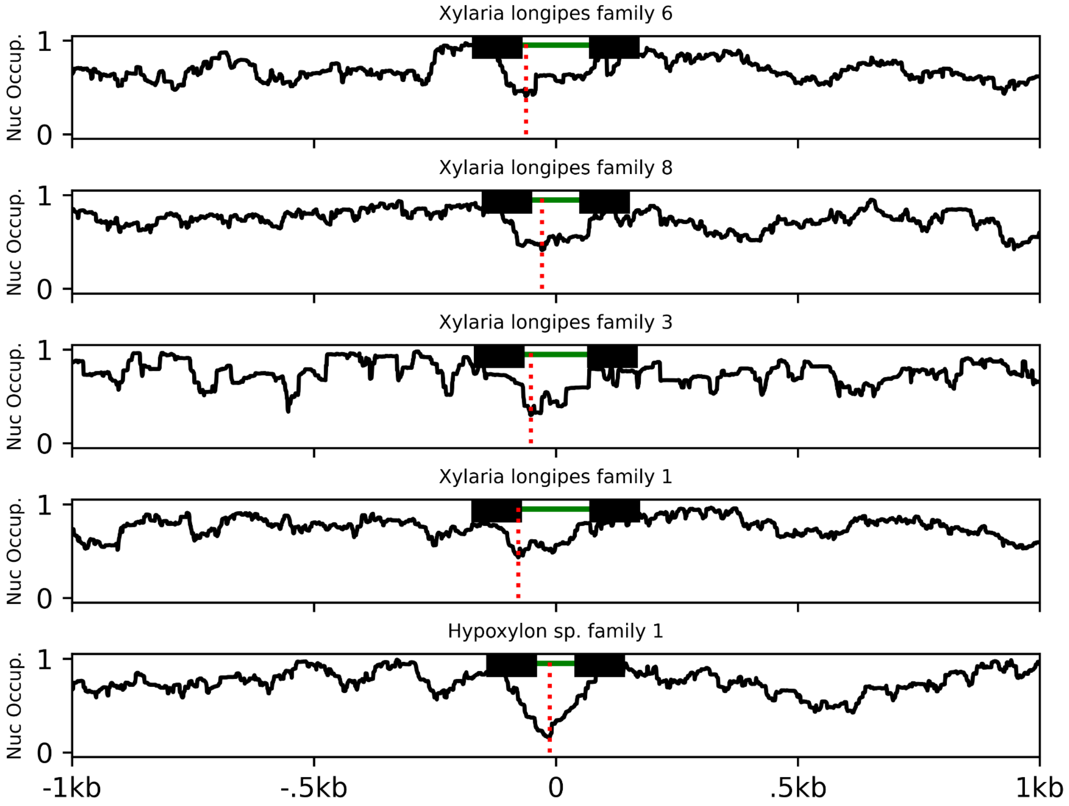
Xylaria longipes family 6
Xylaria longipes family 8
Xylaria longipes family 3
Xylaria longipes family 1
Hypoxylon sp. family 1
Nuc Occup.
1
0
-1kb
-.5kb
0
.5kb
1kb

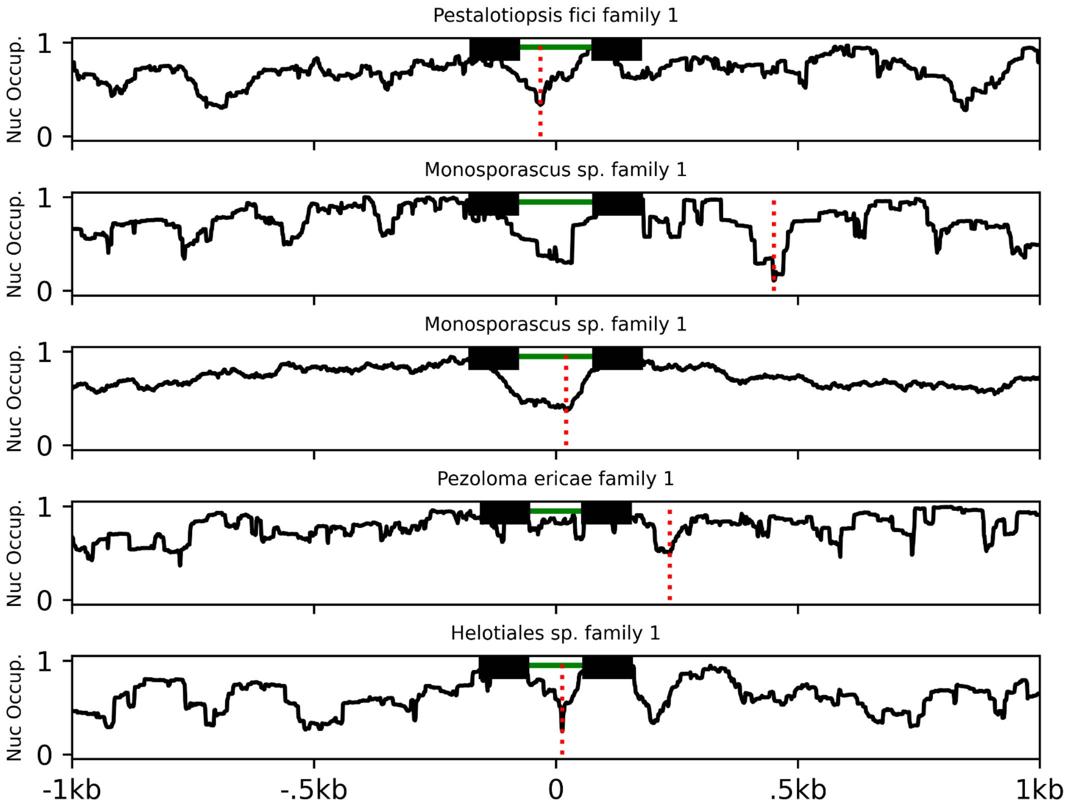
Pestalotiopsis fici family 1
Monosporascus sp. family 1
Monosporascus sp. family 1
Pezoloma ericae family 1
Helotiales sp. family 1
Nuc Occup.
1
0
-1kb
-.5kb
0
.5kb
1kb

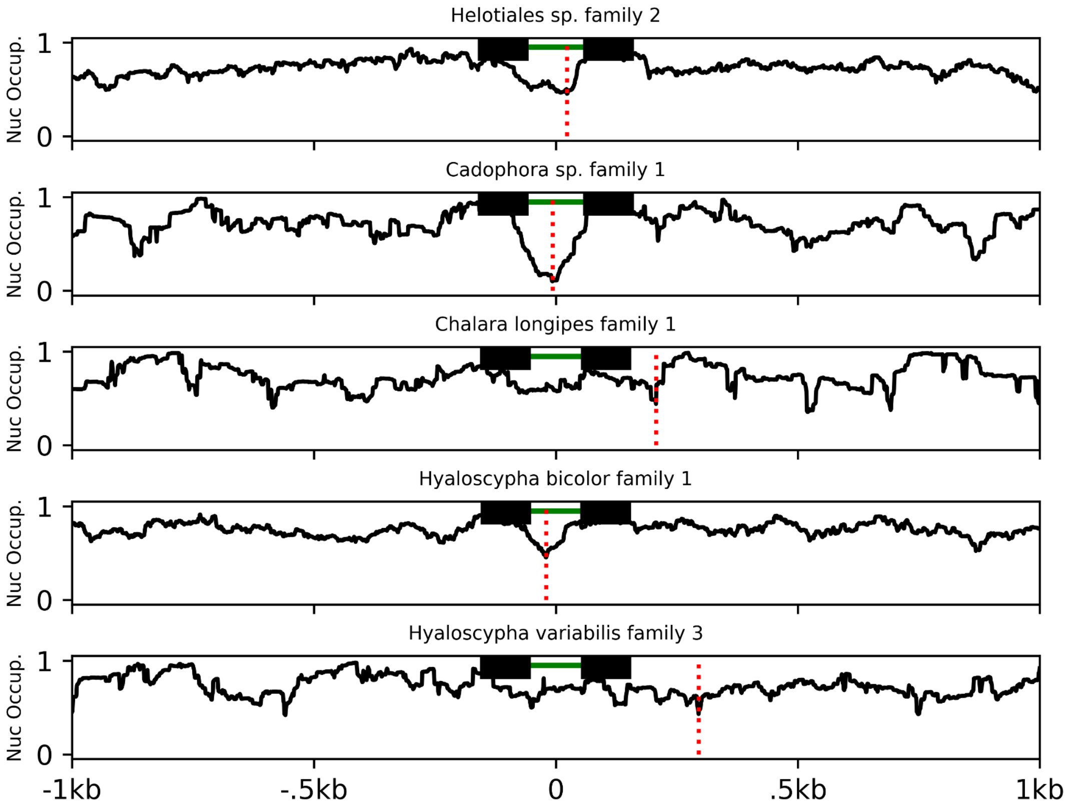

Helotiales sp. family 2
Cadophora sp. family 1
Chalara longipes family 1
Hyaloscypha bicolor family 1
Hyaloscypha variabilis family 3
Nuc Occup.
1
0
-1kb
-.5kb
0
.5kb
1kb

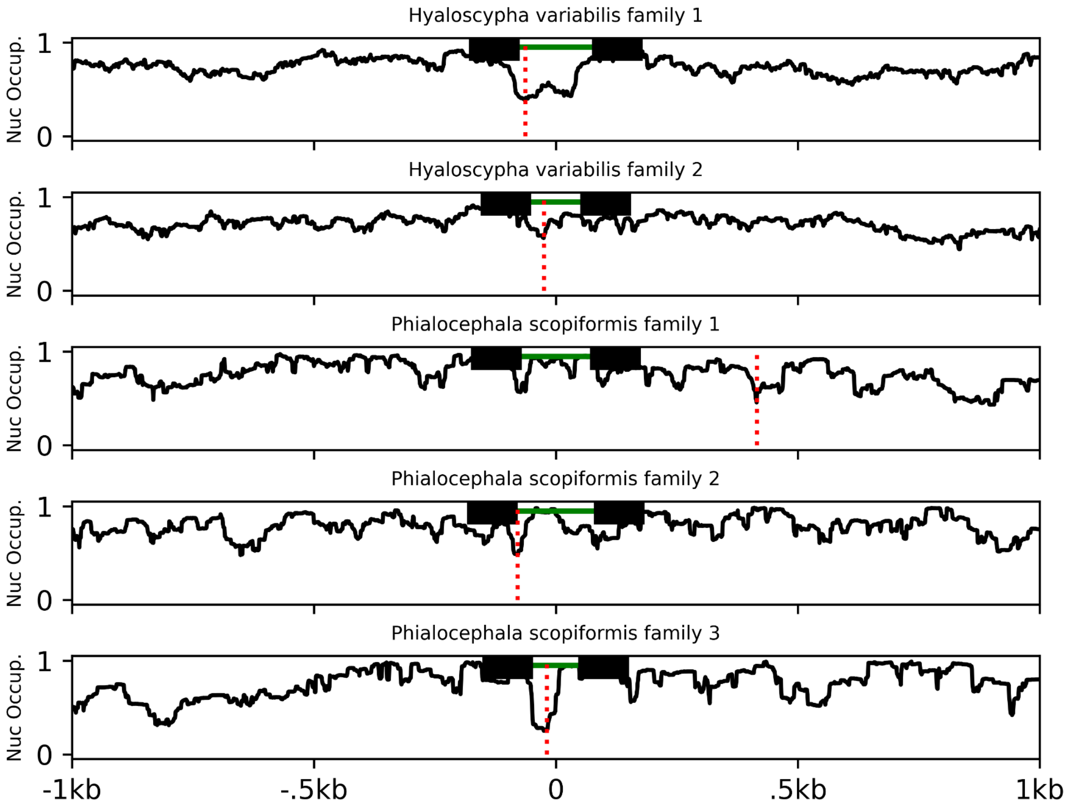
Hyaloscypha variabilis family 1
Hyaloscypha variabilis family 2
Phialocephala scopiformis family 1
Phialocephala scopiformis family 2
Phialocephala scopiformis family 3
Nuc Occup.
1
0
-1kb
-.5kb
0
.5kb
1kb

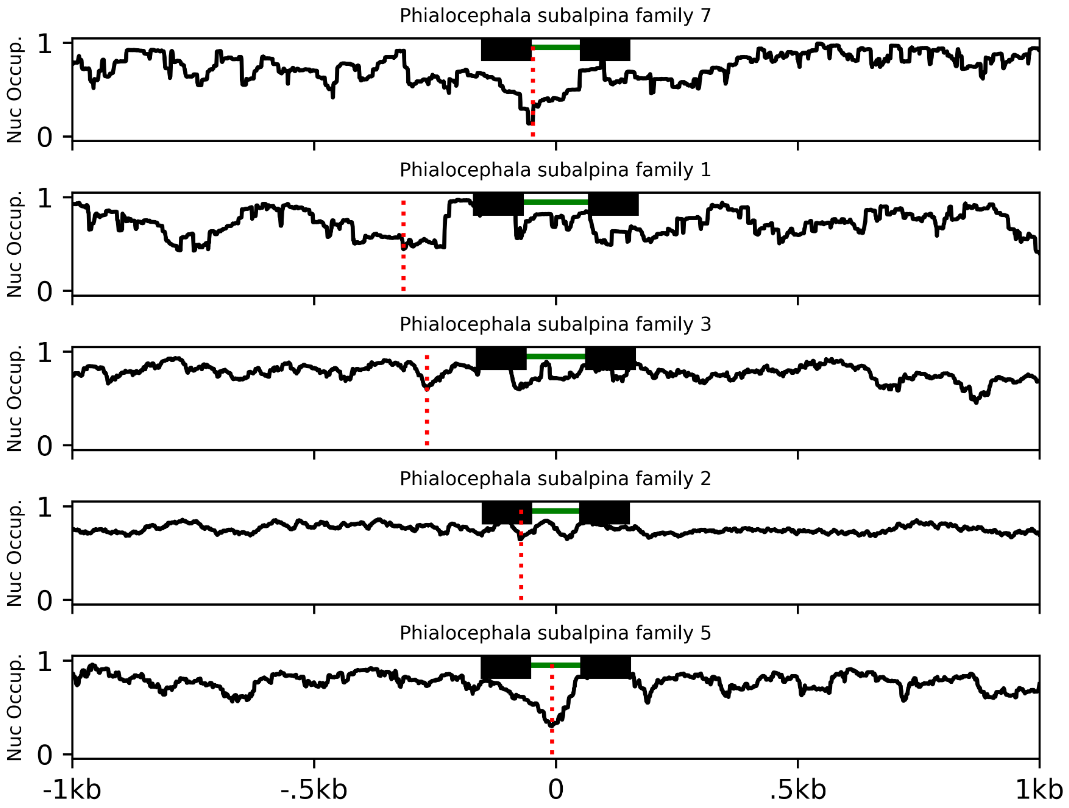
Phialocephala subalpina family 7
Phialocephala subalpina family 1
Phialocephala subalpina family 3
Phialocephala subalpina family 2
Phialocephala subalpina family 5
Nuc Occup.
1
0
-1kb
-.5kb
0
.5kb
1kb

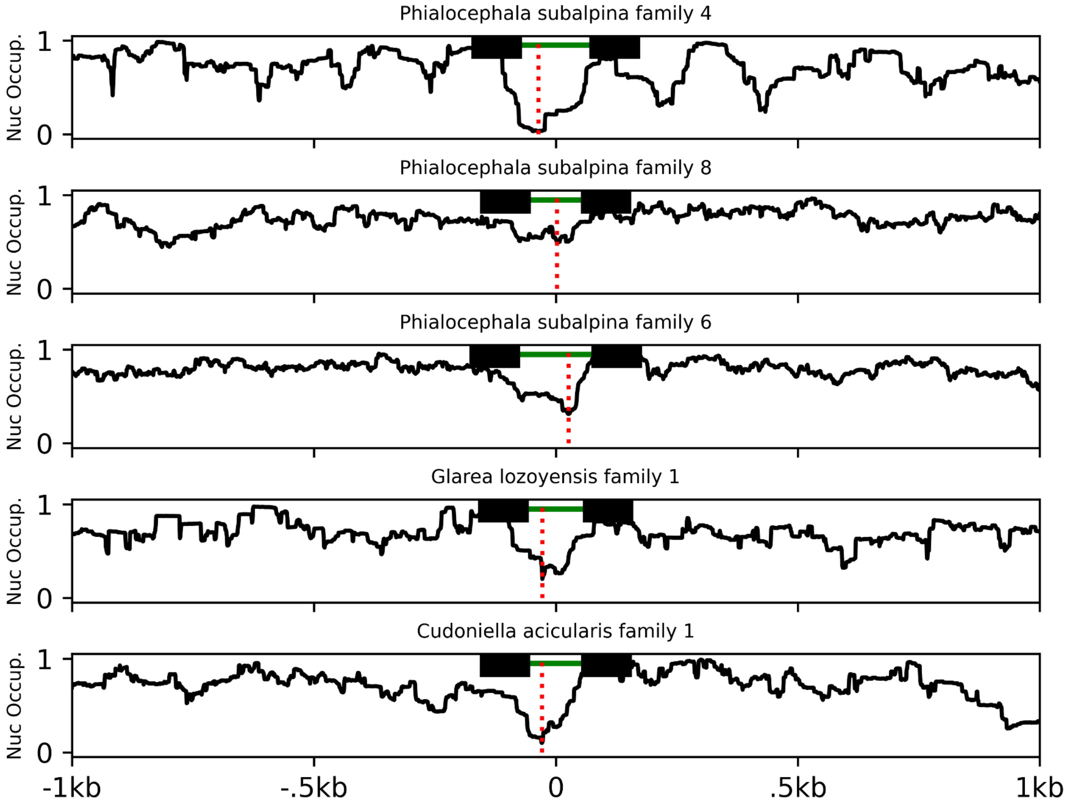
Phialocephala subalpina family 4
Phialocephala subalpina family 8
Phialocephala subalpina family 6
Glarea lozoyensis family 1
Cudoniella acicularis family 1
Nuc Occup.
1
0
-1kb
-.5kb
0
.5kb
1kb

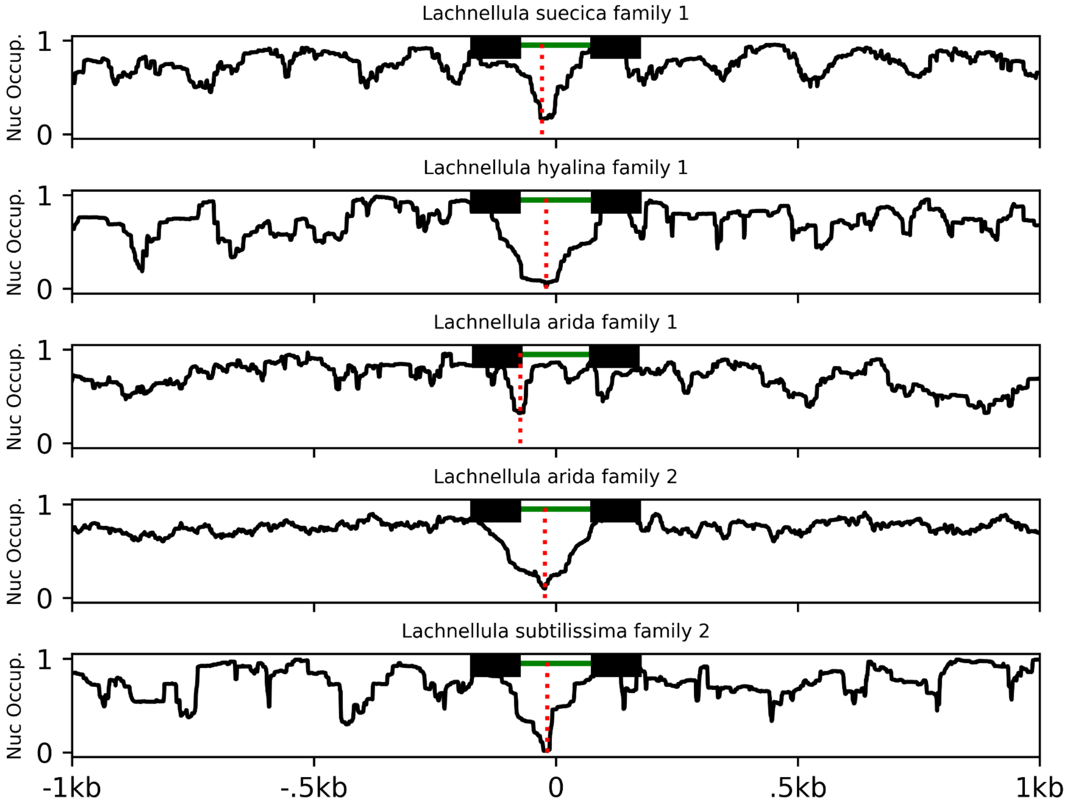


**Supplementary Figure 3: Average nucleosome occupancy relative to Introner position across all families in each species.** Line plots show the average nucleosome occupancy for all Introners for each family on which we could make nucleosome predictions relative to Introner position. The green bar at the top of each plot represents the Introner and lack boxes represent neighboring exons. The red dotted line represents the position at which we observe the minimum nucleosome occupancy in the given region. Coordinates are shown with reference to Introner position. We find that the lowest trough in nucleosome occupancy for the 2kb region surrounding and containing Introners often exists either near or within Introners in most families, further suggesting that Introners inhabit nucleosome linker regions. In some Introner families, such as those found in Blastocystis sp., predictions are noisy due to either a small sample size on which to make predictions (small families of short Introners) or sequence features inhibiting accurate predictions related to specific species (see section on nucleosome prediction above).

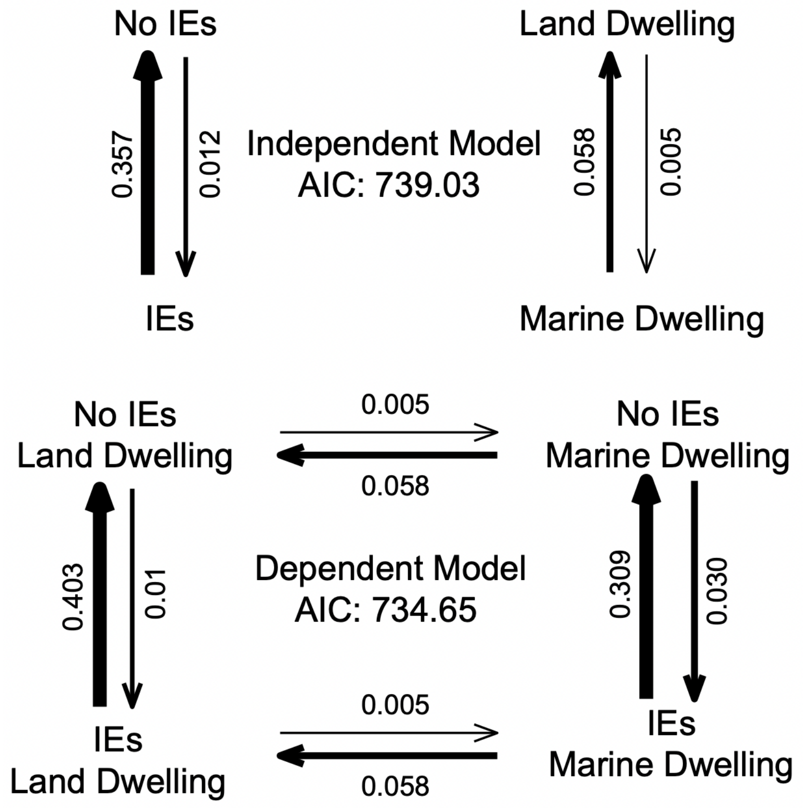


**Supplementary Figure 4: The model in which Introner evolution is dependent on germline accessibility is a better fit to our data than a model in which the two traits evolve independently.** We used Pagel's test to calculate the transition rates for four models (independent, Introner presence dependent on germline accessibility, germline accessibility dependent on Introner presence, and co-dependent) with the former two shown here. Based on the AIC, the dependent model better fits our data ($P < 4.1 \times 10^{-4}$).

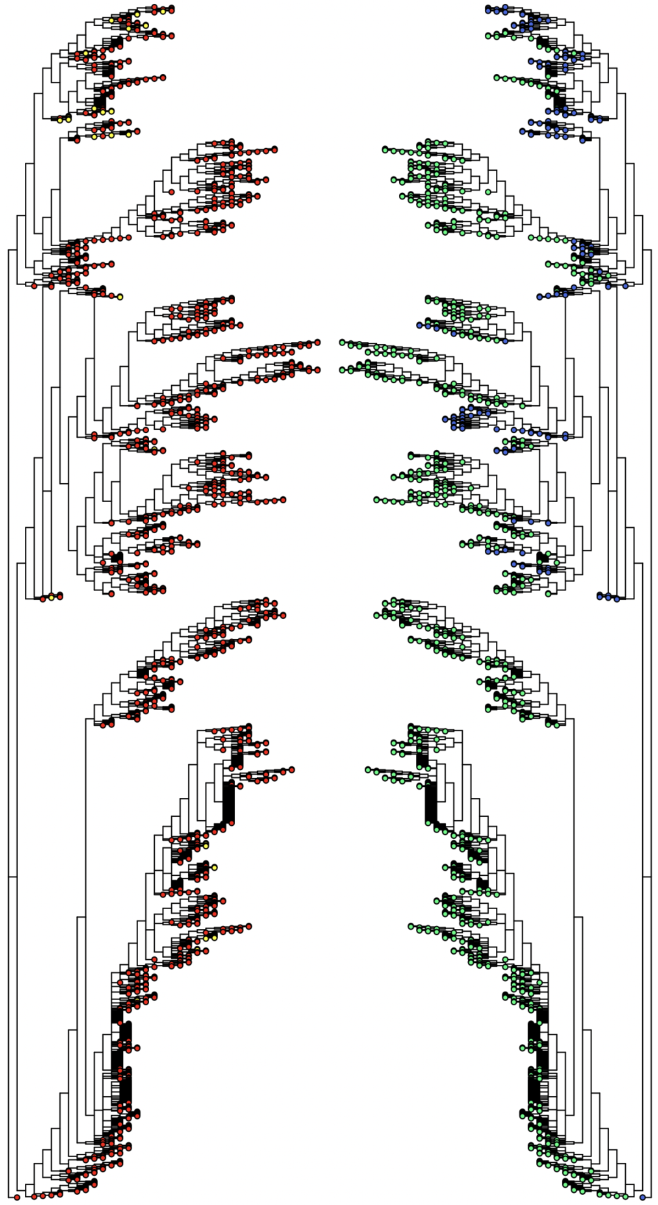

**Supplementary Figure 5: Tree for association with aquatic species.** The phylogeny used to test for association between the presence of Introners and aquatic organisms is shown with marine dwelling species in blue and land dwelling species in green (right), and with Introner-containing species in yellow and those without Introners in red (left). The phylogeny is mirrored for comparisons between the two sets of traits.

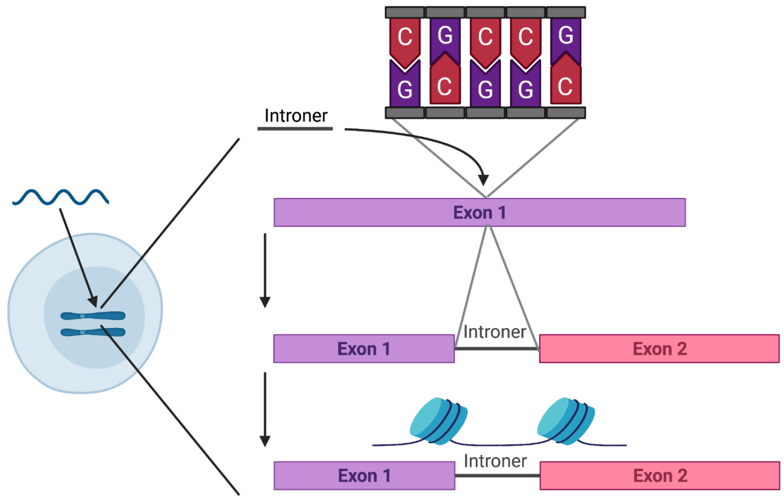


**Supplementary Figure 6: Model for intron gain.** Foreign DNA containing an Introner enters a species (in this case a unicellular eukaryotic organism) via horizontal gene transfer. The Introner transposes into the new host's germline, inserting preferentially into GC rich regions and causing intron gain upon insertion into an exon. Introners generally inhabit nucleosome linker regions whether they preferentially insert in linker regions or cause a change in nucleosome profiles after insertion.
